# New from old: discovery of the novel antibiotic actinomycin L in *Streptomyces* sp. MBT27

**DOI:** 10.1101/2021.10.12.464064

**Authors:** Nataliia Machushynets, Somayah S. Elsayed, Chao Du, Maxime A. Siegler, Mercedes de la Cruz, Olga Genilloud, Thomas Hankemeier, Gilles P. van Wezel

## Abstract

Streptomycetes are major producers of bioactive natural products, including the majority of the antibiotics. While much if the low-hanging fruit has been discovered, it is predited that less than 5% of the chemical space has been mined. Here, we describe the novel actinomycins L_1_ and L_2_, which are produced by *Streptomyces* sp. MBT27. The molecules were discovered via metabolic analysis combined with molecular networking of cultures grown with different combinations of carbon sources. Actinomycins L_1_ and L_2_ are diastereoisomers, and the structure of actinomycin L_2_ was resolved using NMR and single crystal X-ray crystallography. Actinomycin L is formed via a unique spirolinkage of anthranilamide to the 4-oxoproline moiety of actinomycin X_2,_ prior to the condensation of the actinomycin halves. Feeding anthranilamide to cultures of *Streptomyces antibioticus*, which has the same biosynthetic gene cluster as *Streptomyces* sp. MBT27 but only produces actinomycin X_2_, resulted in the production of actinomycin L. This shows that actinomycin L results from joining two distinct metabolic pathways, namely those for actinomycin X_2_ and for anthranilamide. Actinomycins L_1_ and L_2_ showed significant antimicrobial activity against Gram-positive bacteria. Our work shows how new molecules can still be identified even in the oldest of natural product families.

**IMPORTANCE:** Actinomycin was the first antibiotic discovered in an actinobacterium by Selman Waksman and colleagues, as early as 1940. This period essentially marks the start of the ‘golden era’ of antibiotic discovery. Over time, emerging antimicrobial resistance (AMR) and the declining success rate of antibiotic discovery resulted in the current antibiotic crisis. We surprisingly discovered that under some growth conditions, *Streptomyces* sp. MBT27 can produce actinomycins that are significantly different from those that have been published so far. The impact of this work is not only that we have discovered a novel molecule with very interesting chemical modifications in one of the oldest antibiotics ever described, but also that this requires the combined action of primary and secondary metabolic pathways, namely the biosynthesis of anthranilamide and of actinomycin X_2_, respectively. The implication of the discovery is that even the most well-studied families of natural products may still have surprises in store for us.

## INTRODUCTION

Considering the emerging crisis of antibiotic resistance that spreads among bacterial pathogens and increasing incidence of cancer, the search for new, efficient and less toxic drugs remains a priority (46, 56). Actinobacteria have been the source for the majority of the antibiotics in use today (2, 4). Of the Actinobacteria, members of the genus *Streptomyces* produce over half of all currently characterized antibiotics. Genome sequencing revealed that Actinobacteria have much more biosynthetic potential to produce bioactive molecules than originally anticipated, with even the model organisms harbouring many so-called cryptic or silent biosynthetic gene clusters (BGCs) that specify yet unknown compounds (3, 14, 23). Triggering the expression of silent BGCs by genetic and cultivation-based techniques should facilitate unlocking this yet unexplored chemical diversity, allowing the discovery of novel molecules (40, 49). This strategy relies on altering the regulatory networks of the producing organism in response to fluctuating culturing conditions, such as carbon, nitrogen or phosphate concentration (41, 48, 50). Manipulation of fermentation conditions of promising producer strains, known as “one strain many compounds” (OSMAC) approach, is effective in enhancing secondary metabolites production (7, 38). Novel secondary metabolites have been discovered via modification of cultivation parameters, including nutrients (60, 61), and addition of chemical elicitors (11, 37).

Metabolic profiling of crude extracts obtained under different growth conditions represents a challenging analytical task since these mixtures are composed of hundreds of natural products. Therefore, metabolomics, particularly those based on mass spectrometry (MS), became more and more valuable and greatly increased the efficiency of such screenings. Supervised statistical methods are able to classify a response like a biological activity, and to determine the most discriminant metabolite(s) related to such response (57). Moreover, simultaneous dereplication of differentially expressed compounds is implemented into the drug-discovery pipelines in order to avoid rediscovery of already known compounds (18). MS-based metabolomics provides important information on the distribution of the metabolites that are present in complex mixtures, but the identification of their structures is complicated. For this purpose, the Global Natural Products Social Molecular Networking (GNPS) platform was developed, applying both molecular networking and automated searches of tandem mass spectrometry (MS/MS) fragmentation spectra against spectral libraries, to identify structural relationships between metabolites (34, 53). This greatly facilitates the annotation and dereplication of known molecules.

Actinomycin is a DNA-targeting antibiotic and anticancer compound discovered in 1940 by Waksman & Woodruff, and in fact the first antibiotic that was isolated from an actinobacterium (52). Actinomycins are produced by various *Streptomyces* strains and are composed of a chromophore group and two pentapeptide chains with a variable composition of amino acids (25). Actinomycins D, X_0β_ and X_2_ are usually simultaneously produced and differ from each other by substitutions on the “proline” residue in their pentapeptide lactone rings, while members of the actinomycin C complex vary in their “D-valine” residues (12). The pentapeptide precursors are biosynthesized by a non-ribosomal peptide synthetase (NRPS) assembly line, and actinomycins are formed through oxidative condensation of two 3-hydroxy-4-methylanthranilic acid (4-MHA) pentapeptide lactones (PPLs) (13).

In this work we report the discovery of new actinomycin analogues, actinomycin L_1_ and L_2_, from the extracts of *Streptomyces* sp. MBT27. Multivariate data analysis combined with molecular networking indicated that the antimicrobial activity of the extracts correlated with novel actinomycins L_1_ and L_2_ and known actinomycins D, X_0β_ and X_2_. NMR and single crystal X-ray crystallography revealed that an anthranilamide moiety was spiro-linked to a proline residue in the structure of actinomycins L_1_ and L_2_. Such a structural feature has not previously been identified in naturally occurring actinomycins.

## RESULTS

### The influence of carbon sources on bioactivity and actinomycin production

*Streptomyces* sp. MBT27 is a gifted natural product producer that was isolated from Qinling mountains in China, with potent antibacterial activity against various MDR (multidrug-resistant) bacteria (62). We previously showed that the strain among others produces the novel quinazolinones A and B (32). To investigate the antibiotic activity of *Streptomyces* sp. MBT27 the strain was fermented in minimal medium (MM) with either of the following carbon sources (percentages in w/v): 1% of both mannitol and glycerol, 1% mannitol, 2% mannitol, 1% glycerol, 2% glycerol, 1% glucose, 2% glucose, 1% fructose, 1% arabinose, or 1% *N*-acetylglucosamine (GlcNAc). Supernatants of *Streptomyces* sp. MBT27 cultures were extracted with ethyl acetate and bioactivity assays were performed against *Bacillus subtilis* 168. Interestingly, the carbon sources had a huge effect on the antimicrobial activity (Figure S1). Particularly strong antimicrobial activity was observed when the culture medium was supplemented with glycerol + mannitol, glucose 1%, glycerol, fructose or GlcNAc; as compared to when mannitol or arabinose were used as the carbon sources.

In order to investigate the metabolic differences due to nutritional supplementation and correlate that to the antimicrobial activity, LC-MS-based metabolomics was performed. Initially, the LC-MS data were explored by unsupervised Principal Component Analysis (PCA). The first two PCs accounted for 37% and 16%, respectively, of the total data variation. PCA analysis failed to show significant metabolic separation in relation to the observed bioactivity (Figure 1A). The supervised Orthogonal Partial Least Squares Discriminant Analysis (OPLS-DA) was then applied to discriminate the samples based on their ability to inhibit *B. subtilis* (Figure 1B). The cross-validation metrics of the model (*R*^*2*^*Y* = 0.748 and *Q*^*2*^*Y* = 0.676) indicated that the model has a good reliability and ability of prediction. A permutation test was performed (*n* = 100) and the resulting *R*^*2*^*Y* and *Q*^*2*^*Y* values were significantly lower (*p* values < 0.01 for both), which indicated that there was no overfitting in the model (10) (Figure S2). The OPLS-DA loadings S-plot revealed the most discriminative features between active and inactive groups (Figure 1C).

**Figure 1.**
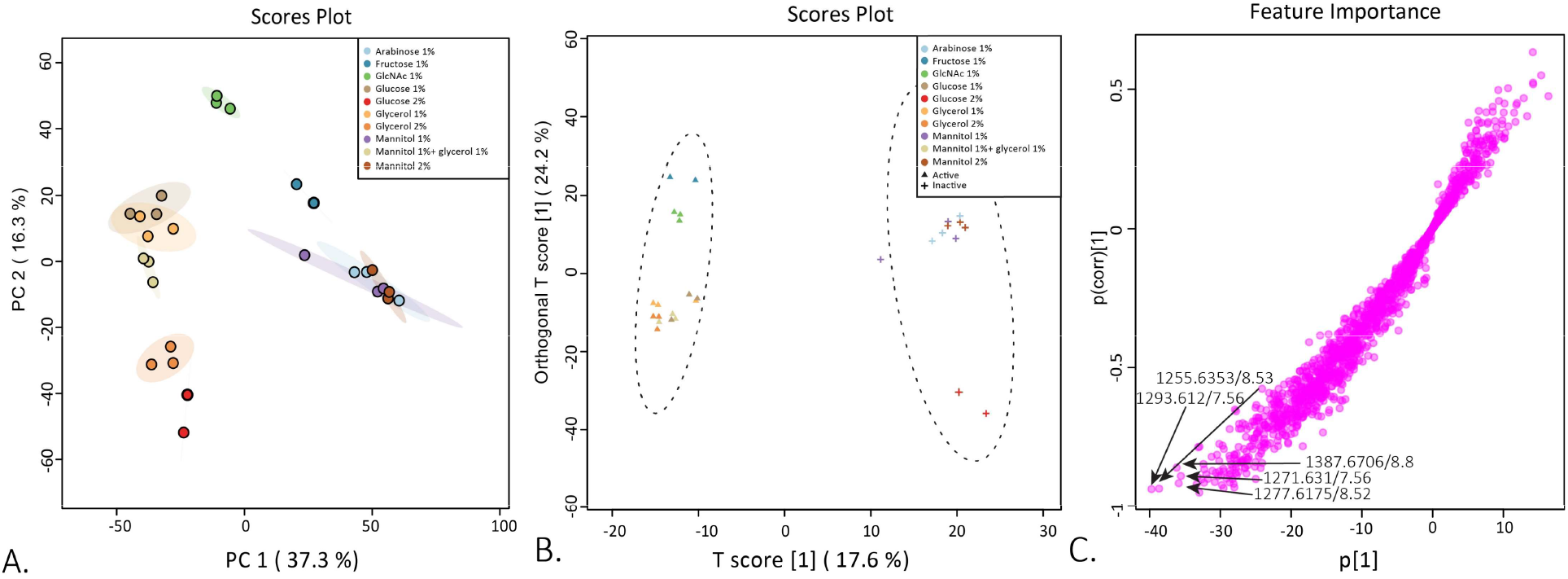
Differential production of metabolites depending on the carbon source. **A**. PCA score plot of *Streptomyces* sp. MBT27 metabolites produced in cultures with different carbon sources, namely, 1% arabinose, 1% fructose, 1% GlcNAc, 1% glucose, 2% glucose, 1% glycerol, 2% glycerol, 1% mannitol, 1% mannitol + 1% glycerol and 2% mannitol (%ages in w/v). **B**. OPLS-DA score plot. Triangles and crosses represent samples of active and inactive groups respectively, circular areas represent the 95% confidence region of each group. **C**. OPLS-DA loadings S-plot. Arrows indicate the most discriminative features that positively correlate with the active groups.

The mass features that correlated best to the bioactivity were *m/z* 1387.6706 (8.8 min), *m/z* 1255.6353 (8.53 min), *m/z* 1277.6175 (8.52 min), *m/z* 1271.651 (7.56 min) and *m/z* 1293.612 (7.56 min) (Figure 1C). Dereplication of those features was performed through comparison of the UV spectra, accurate masses, isotope distribution and fragmentation patterns obtained in MS/MS analysis against the chemistry databases Reaxys, ChemSpider, and the microbial natural products database Antibase (30). This allowed us to annotate the mass features with *m/z* 1255.6353 and 1277.6175 as the [M+H]^+^ and [M+Na]^+^ adduct ions of actinomycin D, respectively, while the mass features with *m/z* 1271.651 and 1293.612 were annotated as [M+H]^+^ and [M+Na]^+^ adduct ions of actinomycin X_0β_, respectively (13). However, the mass feature with an *m/z* value of 1387.6706 [M+H]^+^ could not be matched to any of the previously reported microbial natural products.

Global Natural Product Social (GNPS) molecular networking (53) was subsequently employed to detect MS/MS-based structural relatedness among molecules in an automated manner. The web-based platform generates a molecular network wherein molecules with related scaffolds cluster together. Cytoscape 3.7.2 was used for visualization of the generated molecular networks (45). A network representing the ions detected in the crude extract of *Streptomyces sp*. MBT27 grown with 1% glycerol was constructed, revealing 172 nodes clustered in 10 spectral families (Figure 2). The molecular network revealed an actinomycin spectral family containing actinomycin D, X_2_ and X_0β_. Moreover, the same spectral family included a yet unidentified compound with *m/z* 1387.67. It was closely connected (cosine score > 0.7) to the known actinomycins, suggesting that the molecule was a novel actinomycin. Statistical analysis showed that the extracts with stronger antimicrobial activity contained higher concentrations of actinomycins X_2,_ X_0β_, D and the new compound, in comparison with the less active ones (ANOVA, *p* < 0.05; Figure 2). It is important to note that actinomycins were only detected in the bioactive fractions.

**Figure 2.**
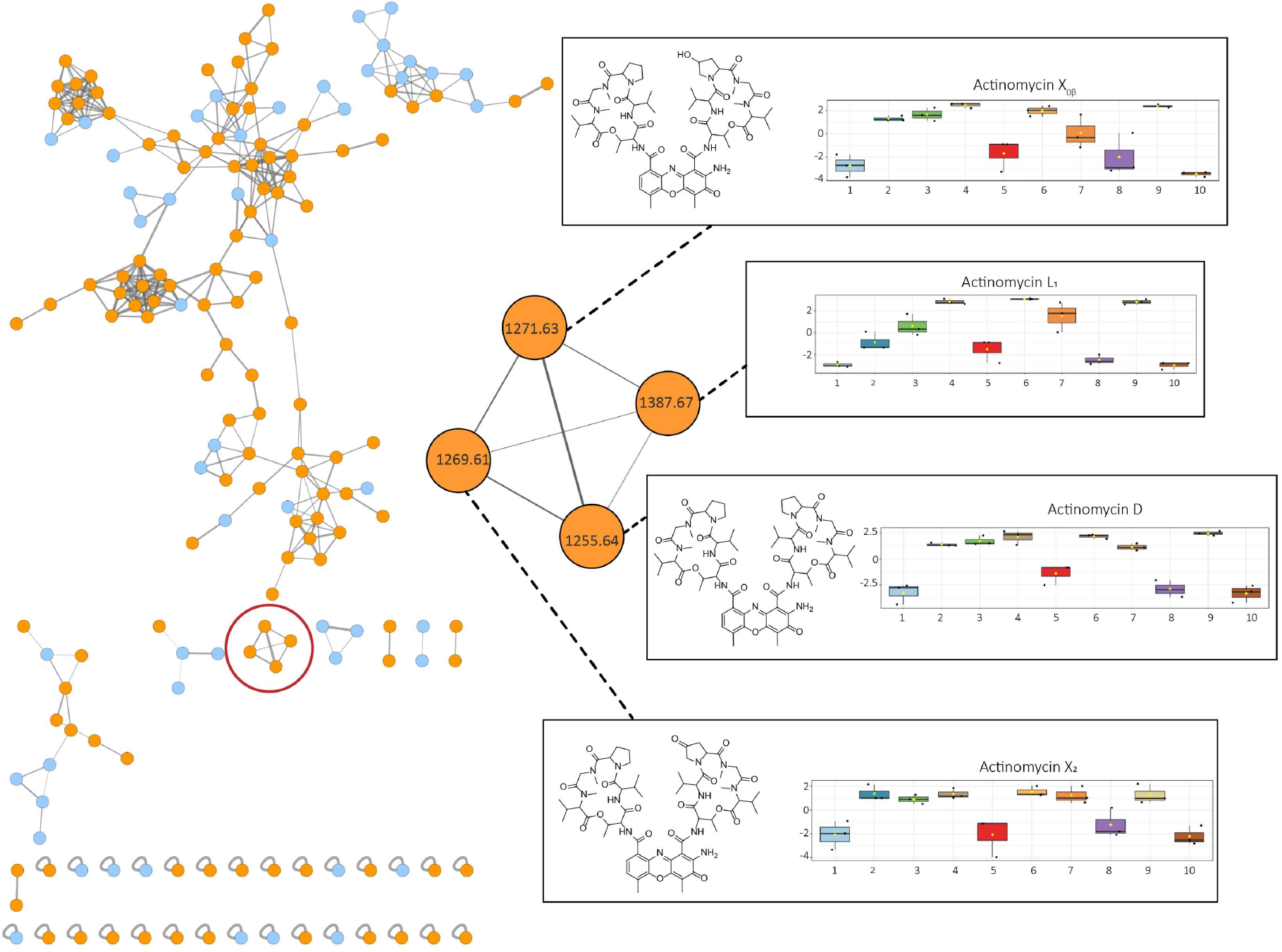
GNPS molecular network of the ions detected in the crude extract of *Streptomyces* sp. MBT27. Cultures were grown for seven days in MM with 1% glycerol. Orange nodes represent ions of the metabolites produced by *Streptomyces* sp. MBT27, while blue nodes represent those of the media components. The actinomycin spectral family is enlarged. Results of ANOVA statistical analysis were mapped onto the molecular network to illustrate the differential production of actinomycin cluster members under various growth conditions. Box plots represent relative intensities of actinomycins X_2_, X_0β_, and D; together with a compound with an *m/z* value of 1387.67, in cultures grown in MM with the following carbon sources: 1. 1% arabinose; 2. 1% fructose; 3. 1% GlcNAc; 4. 1% glucose; 5. 2% glucose; 6. 1% glycerol; 7. 2% glycerol; 8. 1% mannitol; 9. 1% mannitol + 1% glycerol; 10. 2% mannitol (%ages in w/v).

### Large scale fermentation and NMR

To allow identification and structural analysis of the likely novel actinomycin analogue, we performed large-scale fermentation of *Streptomyces* sp. MBT27 followed by bioactivity guided fractionation. The purification process resulted in the isolation of two compounds (**1**) and (**2**), with the same mass (Figure S3 & S4). The NMR spectra of the two compounds were very similar, suggesting that they were diastereoisomers (Figure S5-S11). Based on 1D and 2D NMR analysis of **1**, together with the molecular formula and degrees of unsaturation dictated by the accurate mass, the structure of the isolated diastereoisomers was determined as a variant of actinomycin D, whereby an anthranilamide moiety had been attached through an aminal bond to the γ carbon of one of the proline residues, (Figure 3). The prolyl substitution position is the same as that of the hydroxyl and keto groups in actinomycins X_0β_ and X_2_, respectively. The new actinomycin analogue was designated actinomycin L (with L standing for Leiden, the city of its discovery).

**Figure 3.**
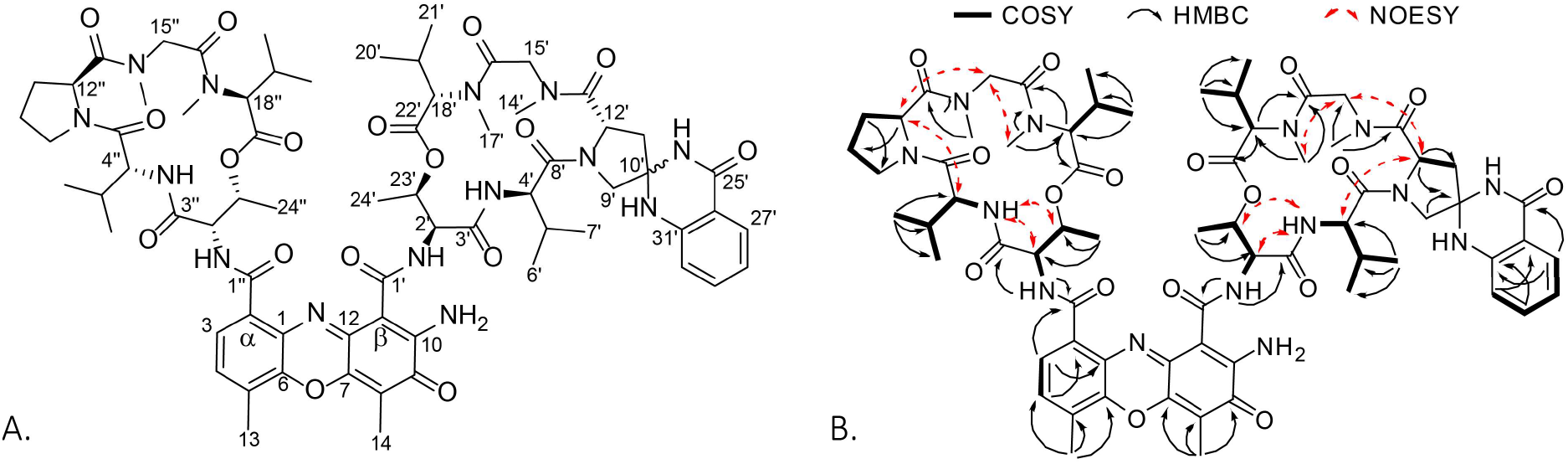
Chemical structures of the new actinomycins. Shown are actinomycin L_1_ (10′*S*) (**1**) and L_2_ (10′*R*) (**2**) (A) and the key COSY, HMBC and NOESY correlations for **1** and **2** (B).

One of the isomers (**2**) was crystallized successfully. Single-crystal X-ray diffraction (Table S1) confirmed the structure obtained from NMR, and established the absolute configuration to be 2′*S*, 2″*S*, 4′*R*, 4″*R*, 10′*R*, 12′*S*, 12″*S*, 18′*S*, 18″*S*, 23′*R*, 23″*R* by anomalous-dispersion effects in diffraction measurements on the crystal (Figure 4). As the absolute configuration of the amino acid residues in **2** was consistent with that of previously reported actinomycins (54), and considering that the two isomers stemmed from the aminal formation at C-10′, compound **1** is inevitably the 10′*S* isomer of actinomycin L.

**Figure 4.**
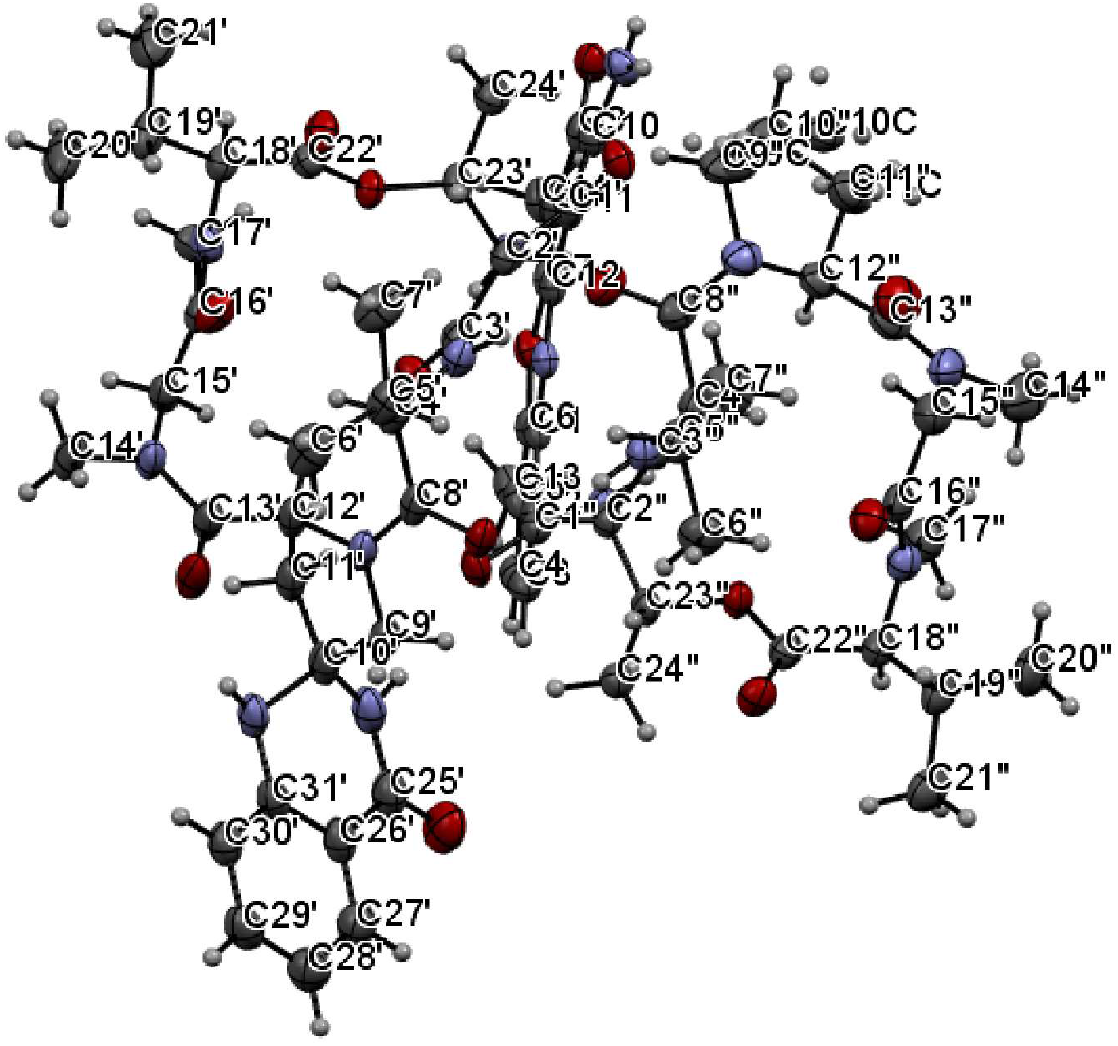
X-ray ORTEP drawing of the crystal structure of compound 2.

### Biosynthesis of actinomycin L

Actinomycins D (or X_1_), X_2_, and X_0β_ detected in the extracts of *Streptomyces* sp. MBT27 are members of the actinomycin X complex. Recently it was shown that actinomycins X_0β_ and X_2_ are formed through the sequential oxidation of the γ-prolyl carbon by the cytochrome P450 enzyme saAcmM (31, 43). Based on its structure, actinomycin L is most likely formed through an aminalization reaction between the two amino groups of anthranilamide and the γ-keto group on the proline residue of actinomycin X_2_. Accordingly, its production should be arrested when one of the precursors is not available. Interestingly, *Streptomyces* sp. MBT27 produced actinomycin L in very low amounts when grown with fructose (1% w/v) as the sole carbon source. Moreover, ANOVA statistical analysis showed that anthranilamide was produced in equally low amounts under the same growth conditions (ANOVA, *p* < 0.05; Figure 5). Under conditions where *Streptomyces* sp. MBT27 produced actinomycin L, namely when grown in MM with 1% GlcNAc, 1% glucose, 1% glycerol, 2% glycerol or in 1% mannitol + 1% glycerol, the strain invariably produced both actinomycin X_2_ and anthranilamide. However, under conditions where actinomycin X_2_ was produced but not anthranilamide, the strain failed to produce actinomycin L (Figure 5).

**Figure 5.**
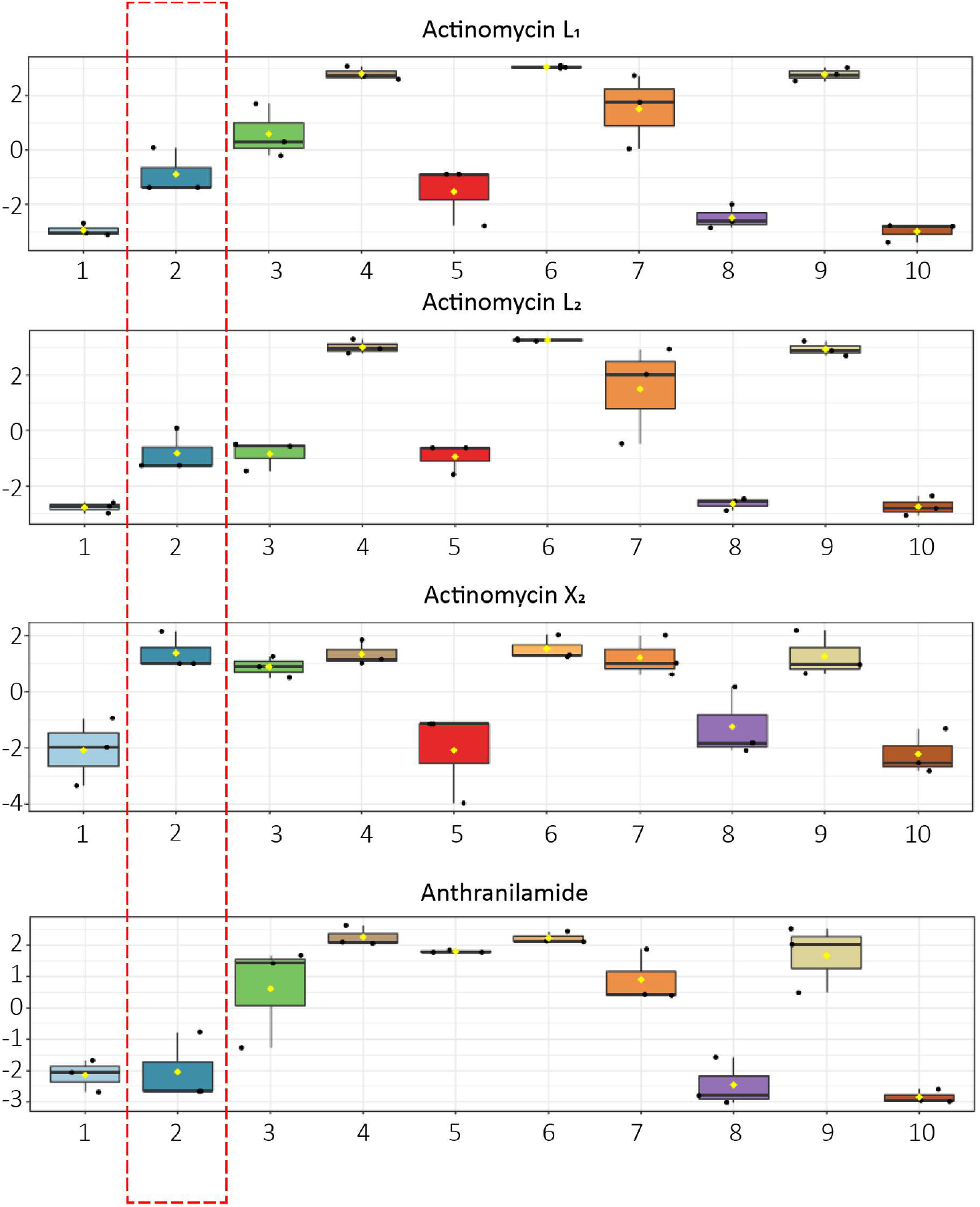
Box plots showing the relative intensities of actinomycin L_1_, L_2_ and X_2_ and anthranilamide. Cultures of *Streptomyces* sp. MBT27 were grown for seven days in MM with the following carbon sources: 1. 1% arabinose; 2. 1% fructose; 3. 1% GlcNAc; 4. 1% glucose; 5. 2% glucose; 6. 1% glycerol; 7. 2% glycerol; 8. 1% mannitol; 9. 1% mannitol +1 % glycerol; 10. 2% mannitol (% ages in w/v). Red box indicates the abundance of actinomycin L_1_, L_2_ and X_2_ and anthranilamide in the cultures grown with fructose (1% w/v). Note that *Streptomyces* sp. MBT27 produced actinomycin L and anthranilamide in very low amounts when fermented with fructose (1% w/v) as the sole carbon source.

We therefore wondered if anthranilamide may be a precursor for the biosynthesis of actinomycin L. To test this hypothesis, we performed a feeding experiment, whereby anthranilamide was added to cultures of *Streptomyces* sp. MBT27 grown in MM with 1% fructose, where virtually no actinomycin L was produced. Analysis of the supernatant of the cultures via LC- MS revealed that actinomycin L was readily produced when anthranilamide was added, but not without it (Figure 6A). This strongly suggested that anthranilamide is required for the production of actinomycin L. However, extracts of *Streptomyces* sp. MBT27 fermented with 1% fructose and additional anthranilic acid contained both anthranilamide and actinomycin L (Figure 6A). This suggests that indeed anthranilic acid is converted into anthranilamide, which in turn is incorporated into actinomycin L.

**Figure 6.**
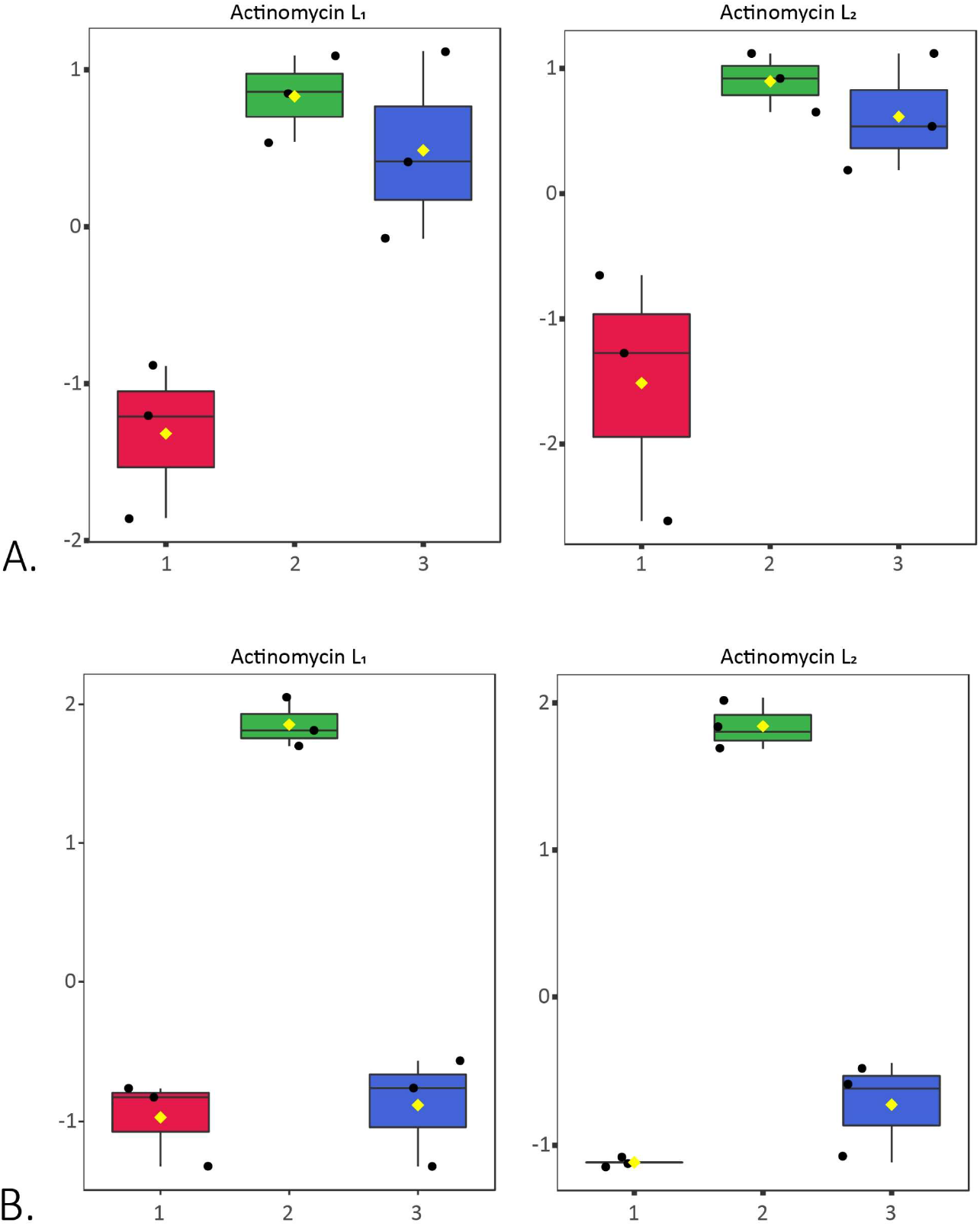
Anthranilamide is required for the biosynthesis of actinomycins L_1_ and L_2_. Box plots show the relative intensities of actinomycin L_1_ and L_2_ in the cultures of *Streptomyces* sp. MBT27 (A) and *S. antibioticus* (B) fermented for seven days in MM with fructose (1% w/v) (1), fed with 0.7 mM anthranilamide (2) and 0.7 mM anthranilic acid (3). Note that *S. antibioticus* produces actinomycin L exclusively in the presence of anthranilamide and not with anthranilic acid; conversely, *Streptomyces* sp. MBT27 is able to convert anthranilic acid into anthranilamide, enabling the production of actinomycin L.

In order to unambiguously verify that actinomycin L was the product of anthranilamide and actinomycin X_2_, we conducted another biotransformation experiment, now feeding anthranilamide to *S. antibioticus* IMRU 3720, which is a known producer of actinomycins X_2_ and X_0β_, but fails to produce actinomycin L under any condition tested. In line with our hypothesis, *S. antibioticus* IMRU 3720 also failed to produce anthranilamide under any of the growth conditions (Figure S16). Excitingly, LC-MS analysis revealed the production of actinomycin L by *S. antibioticus* IMRU 3720 when anthranilamide was fed to the cultures, but never without anthranilamide (Figure 6B). This validates the concept that anthranilamide is a key precursor of actinomycin L. Conversely, when anthranilic acid instead of anthranilamide was added to cultures of *S. antibioticus* IMRU 3720, we failed to detect actinomycin L and anthranilamide (Figure 6B).

The oxidation of the proline residue in actinomycins X_0β_ and X_2_ occurs following the formation of the two halves of actinomycin, known as 4-MHA PPLs, and prior to the condensation of these halves to form actinomycin (43). Taking this into account we reasoned that anthranilamide should be incorporated into the actinomycin halves prior to condensation. To check this, 3- hydroxy-4-methylbenzoic acid (4-MHB) was added to cultures of *Streptomyces* sp. MBT27 and of *S. antibioticus* IMRU 3720. 4-MHB is a structural analogue of 4-MHA that replaces 4-MHA as a starter unit in the nonribosomal assembly of the actinomycin halves (43). When 4-MHB replaces 4-MHA, 4-MHB containing PPLs accumulate, because they cannot react with each other to give a phenoxazinone ring, as is the case with 4-MHA PPLs (27). LC-MS analysis of the 4-MHB- supplemented extracts showed the appearance of the previously reported 4-MHB-containing pentapeptide lactones PPL 1, PPL 0, and PPL 2, and new PPL, designated as PPL 3 (Figure S17- 20, Table S2). The exact mass and fragmentation pattern of PPL 3 was consistent with a 4-MHB containing PPL wherein an anthranilamide moiety had been attached to the proline residue (Figure S20).

Taken together, the feeding experiments convincingly show that actinomycin L is formed through reaction of anthranilamide with the 4-keto group on the proline residue in the pentapeptide lactone. Moreover, results of the feeding experiments with 3-hydroxy-4-methylbenzoic acid show that this reaction occurs prior to the condensation of the pentapeptide lactones into actinomycin L (Figure 7).

**Figure 7.**
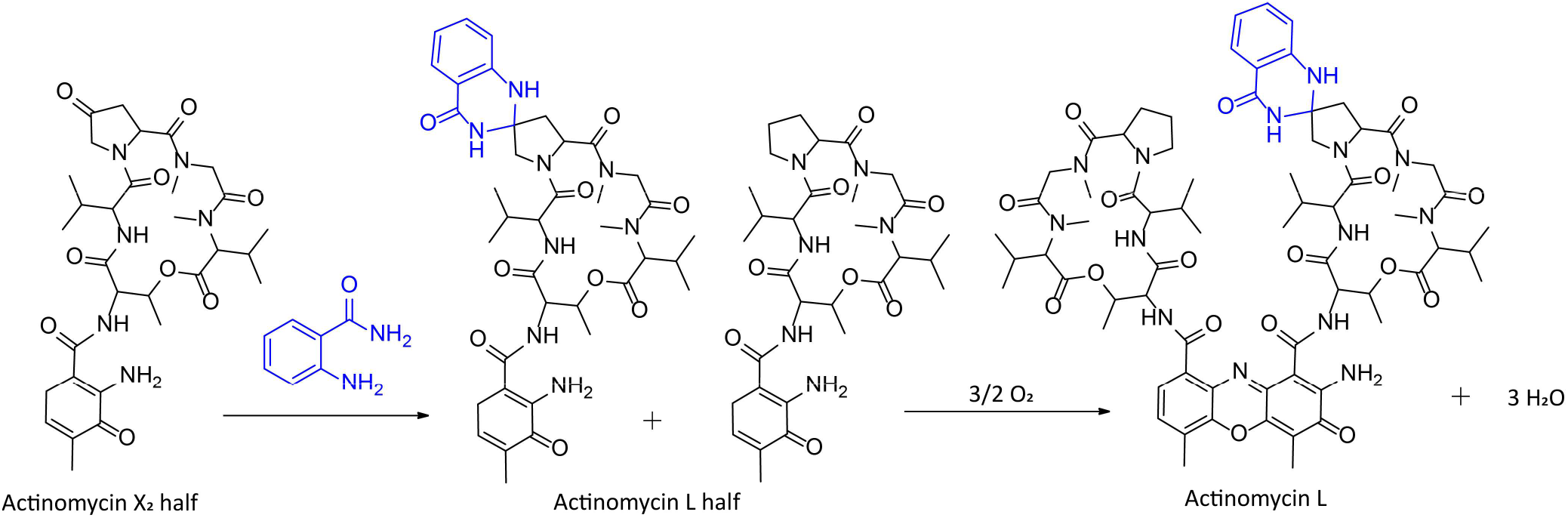
Proposed biosynthetic pathway for actinomycin L. We propose that actinomycin L is formed through the reaction of anthranilamide (blue) with the 4-oxoproline moiety of actinomycin X_2_ prior to the condensation of two 4-MHA PPLs into the actinomycin L.

### Identification of the actinomycin BGC in *Streptomyces* sp. MBT27

To characterize the BGC responsible for actinomycin biosynthesis and compare the genes with those found in known actinomycin BGCs, *Streptomyces* sp. MBT27 was sequenced using the PacBio platform. Assembly of the PacBio reads resulted in two contigs of 8.4 Mb and 0.13 Mb in length. Analysis using AntiSMASH 6 (6) readily identified the actinomycin BGC in the 8.4 Mb contig. Comparison to the actinomycin X_2_ BGC from *S. antibioticus* showed that all genes were highly conserved between the two clusters (Table S3 and Fig. S21). This strongly suggests that the actinomycin BGC does not specify the observed modifications in the actinomycin structure, and is not responsible for the production of anthranilamide. We have not yet identified the enzyme for the predicted conversion of anthranilic acid to anthranilamide.

### Bioactivity of isolated compounds (MIC)

Bioactivity assays were carried out for the actinomycins, to test their ability to act as antibiotics. As expected, the compounds showed selective antibacterial activity against Gram-positive pathogens, and none of the actinomycins presented any activity against *E. coli* ATCC25922 or *K. pneumoniae* ATCC700603 (Table 2). All compounds except actinomycin X_0ß_ showed antibacterial activity against Gram-positive bacteria with MIC values ranging from 1 to 16 μg/mL. Actinomycin L_1_ showed somewhat higher bioactivity than actinomycin L_2_, while both compounds showed slightly higher MICs than actinomycin D and Actinomycin X_2_.

## DISCUSSION

Actinomycin was the first antibiotic identified in Actinobacteria (52). The well-established actinomycin structure is composed of a heterocyclic chromophore and two cyclic PPLs. Biosynthetically, PPL is biosynthesized by an NRPS assembly line with the 4-MHA as the initiating unit (24, 35). 4-MHA is derived from 3-hydroxy-4-methylkynurenine (4-MHK), which is formed by methylation of 3-hydroxykynurenine (3-HK) (12). Our work surprisingly revealed a novel structure within the extensively studied actinomycin family, namely that of actinomycin L, which arises via attachment of an anthranilamide moiety to the γ-carbon of one of the proline residues through aminal formation. ANOVA statistical analysis proved that production of anthranilamide is the limiting factor in the biosynthesis of actinomycin L. Feeding experiments with anthranilamide suggested that actinomycin L is formed through the spontaneous reaction of anthranilamide with the 4-oxoproline site of actinomycin X_2_ prior to the condensation of the two 4-MHA PPLs into actinomycin L. To the best of our knowledge, the attachment of anthranilamide to a 4-oxoproline moiety is a novel observation.

The actinomycin BGC of *Streptomyces* sp. MBT27 harbours the same genes as that of *S. antibioticus*, with high homology between the genes, which strongly suggests that the modification of actinomycin X_2_ to actinomycin L is not encoded by the BGC itself. Indeed, we anticipate that anthranilamide is derived from anthranilic acid in *Streptomyces* sp. MBT27, whereby anthranilic acid in turn is biosynthesized through the shikimate pathway (17). Anthranilic acid is a commonly produced primary metabolite in *Streptomyces*, while anthranilamide is less common (5, 21, 44). The actinomycin X_2_ producer *S. antibioticus* IMRU 3720 fails to convert anthranilic acid into anthranilamide, which explains why actinomycin L was also not detected in the extracts. However, actinomycin L was produced when we feeded cultures of *S. antibioticus* IMRU 3720 with additional anthranilamide, which is fully in line with our proposed biosynthetic pathway. Thus, actinomycin L is an example of a natural product that requires the joining of two separate metabolic pathways, and this is a concept that needs more attention. After all, scientists rely increasingly on heterologous expression and synthetic biology approaches (29), and these will likely fail if genes are required that do not fall within the main BGC.

Carbon source utilization plays a major role in the control of antibiotic production (42, 50). Control of carbon utilization in streptomycetes is largely mediated via glucose kinase, via a mechanism that is still not well understood (39, 51). The pathways for antibiotic production are subject to Glk-dependent and Glk-independent control (20). Indeed, the production of actinomycins by *Streptomyces* spp. is strongly influenced by the carbon source, whereby the preferred carbon source varies from strain to strain (16, 22, 26, 47, 55). D-galactose favors actinomycin production in *Streptomyces antibioticus* over arabinose, xylose, glucose, fructose and rhamnose (26), while glycerol was the optimal carbon source for actinomycin production by *S. antibioticus* Tü 6040 and *S. antibioticus* SR15.4 (47) (22). In the case of *Streptomyces* sp. MBT27, growth on MM with glycerol, GlcNAc, fructose and glucose (all 1% w/v) as sole carbon sources were the best carbon sources to promote the production of actinomycins. However, increasing the glucose concentration to 2% blocked the production of actinomycins. Glucose was previously reported to repress the transcription of the gene for hydroxykynureninase, which is involved in the formation of the main actinomycin precursor 4-MHA (26). Importantly, in our experiments the carbon source not only promoted the overall production levels, but also contributed to the chemical diversity of the actinomycins, including the production of actinomycin L. This coincided with the production of anthranilamide, an essential substrate to form this novel actinomycin variant.

In the 21^st^ century, genome mining and renewed drug discovery efforts have revealed that Actinobacteria may produce many more molecules than was expected (33). What is important to note is that this also applies to well-known families of molecules and in extensively studied model organisms. Examples are the highly rearranged cryptic polyketide lugdunomycin that belongs to the family of angucyclines (59), the new glycopeptide corbomycin with a novel mode of action (15), as well as the discovery of coelimycin (19) and a novel branch of the actinorhodin biosynthetic pathway (58) in the model organism *Streptomyces coelicolor*. The discovery of actinomycin L provides another interesting example that we have not yet exhausted the known part of the chemical space. Indeed, the isolation of these novel actinomycins underlines that the biosynthetic potential of Actinobacteria still has major surprises in store, and that we can expect that new molecules can be discovered even within extensively studied microbes and compound classes.

## MATERIALS AND METHODS

### General Experimental Procedures

Optical rotation was recorded on MCP 100 modular circular polarimeter (Anton Paar). FT-IR spectra were recorded on IRSpirit QATR-S Fourier-transform infrared spectrophotometer (Shimadzu Corporation, Japan). UV measurements were performed using a Shimadzu UV-1700 UV-VIS spectrophotometer (Shimadzu Corporation, Japan). NMR spectra were recorded on a Bruker Ascend 850 MHz NMR spectrometer (Bruker BioSpin GmbH). The LC-ESI-MS analyses were performed using Waters Acquity UPLC system coupled to Agilent 6530 QTOF MS equipped with Agilent Jet Stream ESI source (Agilent Technologies, Inc., Palo Alto, CA, USA). The LC-ESI-MS/MS analysis was conducted using Shimadzu Nexera X2 UHPLC system, with attached PDA, coupled to Shimadzu 9030 QTOF mass spectrometer. HPLC purification was performed on Waters preparative HPLC system comprised of 1525 pump, 2707 autosampler, 2998 PDA detector, and Water fraction collector III. All organic solvents were HPLC or LC-MS grade, depending on the experiment.

### Bacterial strain, growth conditions and metabolite extraction

*Streptomyces* sp. MBT27 was obtained from the Leiden University strain collection and had previously been isolated from the Qingling Mountains, Shanxi province, China (62). Cultures were grown in triplicate in 100 mL Erlenmeyer flasks containing 30 mL of liquid minimal medium (MM) (28), supplemented with various carbon sources, and inoculated with 10 μL of 10^9^/mL spore suspension. The carbon sources (percentages in w/v) were: 1% mannitol + 1% glycerol, 1% mannitol, 2% mannitol, 1% glycerol, 2% glycerol, 1% glucose, 2% glucose, 1% fructose, 1% arabinose or 1% *N*-acetylglucosamine (GlcNAc). The cultures were incubated in a rotary shaker at 30 °C and 220 rpm for seven days. Following fermentation, culture supernatants were extracted with ethyl acetate (EtOAc) and evaporated under reduced pressure. In the series of feeding experiments *Streptomyces* sp. MBT27, *S. antibioticus* IMRU 3720 and *S. chrysomallus* ATCC11523 were fermented in MM supplemented with 1% fructose and 0.7 mM anthranilamide. For the directed biosynthesis of non-natural actinomycin X halves *Streptomyces* sp. MBT27 and *S. antibioticus* IMRU 3720 were grown in MM with 1% w/v fructose, 0.7 mM anthranilamide and 0.7 mM 3-hydroxy-4-methylbenzoic acid (4-MHB) and extracted with EtOAc.

### Genome sequencing, assembly and annotation

*Streptomyces* sp. MBT27 was grown in YEME at 30 °C and 220 rpm for 48 hours. DNA was extracted from *Streptomyces* sp. MBT27 as described (28). DNA quality was verified by agarose gel electrophoresis. PacBio sequencing and assembly was performed by Novogene (UK). Generally, library was prepared using SMRTbell template prep kit (PacBio, USA) according to manufacturer instructions. Sequencing was then performed using PacBio Sequel platform in continuous long reads mode. Assembly was done using Flye (version 2.8.1, 10.1038/s41587-019-0072-8). Biosynthetic gene clusters (BGCs) in this genome were annotated using AntiSMASH 6.0 (6). The actinomycin BGC was then extracted and compared with the same cluster from *S. antibioticus* IMRU 3720 using clinker (version 0.0.20, 10.1093/bioinformatics/btab007) with default settings.

### Up-scale fermentation, extraction and fractionation

For large-scale fermentation, *Streptomyces* sp. MBT27 was grown in eight 2 L Erlenmeyer flasks, each containing 500 mL liquid MM supplemented with 2% w/v glycerol at 30 °C for seven days. The metabolites were extracted from the spun culture media using EtOAc, and the solvent was subsequently evaporated under reduced pressure at 40 °C. The crude extract (1.4 g) was adsorbed onto 1.4 g silica gel (pore size 60 Å, 70–230 mesh, Sigma Aldrich), and loaded on silica column, which was eluted using gradient mixtures of n-hexane, acetone, and MeOH. The fractions eluted with *n*-hexane–acetone (1:1) and acetone were combined, reconstituted in MeOH, and injected into preparative SunFire C_18_ column (10 μm, 100 Å, 19 × 150 mm), which was eluted with a H_2_O–MeOH gradient of 50–100% in 20 min, at a flow rate of 15 mL/min. The fraction containing the actinomycins was collected and further purified on semi-preparative SunFire C_18_ column (5 μm, 100 Å, 10 × 250 mm), run at 3 mL/min, and eluted using a H_2_O–MeOH gradient of 70–100% in 20 min, to yield actinomycins L_1_ (**1**, 2.9 mg), L_2_ (**2**, 1.3 mg), X_2_ (**3**, 1 mg), X_0β_ (**4**, 2.9 mg), and D (**5**, 3.2 mg),.

Actinomycin L_1_ (**1**): red amorphous powder; [α] _D_^25^ −38 (c 0.2, MeOH); UV (MeOH) *λ*_max_ (log ε) 211 (1.957), 312 (0.124), 427.5 (0.152), 438 (0.151) nm; IR v_max_ 3301, 2963, 2921, 2859, 1740, 1662, 1644, 1585, 1521, 1465, 1405, 1300, 1262, 1191, 1097 cm^−1^; ^1^H and ^13^C NMR data, see Table 1; HRESIMS (positive mode) *m/z* 1387.6681 [M + H]^+^ (calcd. for C_69_H_91_N_14_O_17_, 1387.6681). Actinomycin L_2_ (**2**): red amorphous powder; [α] _D_^25^ −54 (c 0.2, MeOH); UV (MeOH) *λ*_max_ (log ε) 226 (2.189), 364 (0.186) nm, 426.5 (0.367) nm; IR v_max_ 3308, 2943, 2929, 2831, 1748, 1662, 1644, 1585, 1448, 1406, 1302, 1262, 1191, 1113 cm^−1^; HRESIMS (positive mode) *m/z* 1387.6674 [M + H]^+^ (calcd. for C_69_H_91_N_14_O_17_, 1387.6681). The X-ray diffraction experiment on a crystal grown from MeOH further confirmed the structure and determined the absolute configuration of compound **2** (Figure 4) (CCDC 2110000).

**Table 1.** ^1^H and ^13^C NMR data of 1 in CDCl_3_ at 298 K^[a]^. [a] ^1^H 850 MHz and ^13^C NMR resonances inferred from HSQC and HMBC spectra.

| Position | δ<sub>c</sub>, type | δ<sub>H</sub>, mult. (J in Hz) | Position | δ<sub>c</sub>, type | δ<sub>H</sub>, mult. (J in Hz) |
| --- | --- | --- | --- | --- | --- |
| 1 | 129.5, C |  | 22' | 167.4, C |  |
| 2 | 130.9, C |  | 23' | 74.5, CH | 5.29, qd (6.3, 3.1) |
| 3 | 125.9 | 7.75, m | 24' | 17.5, CH<sub>3</sub> | 1.23, d (6.3) |
| 4 | 130.0, CH | 7.44, dd (7.6, 0.9) | 25' | 163.1, C |  |
| 5 | 128.5, C |  | 26' | 114.3, C |  |
| 6 | 140.6, C |  | 27' | 128.6, CH | 7.90, dd (7.9, 1.6) |
| 7 | 144.7, C |  | 28' | 118.7, CH | 6.81, dt (0.9, 7.9) |
| 8 | 113.6, C |  | 29' | 133.6, CH | 7.30, dt (1.6, 7.9) |
| 9 | 178.9, C |  | 30' | 113.9, CH | 6.67, dd (7.9, 0.9) |
| 10 | ND |  | 31' | 146.7, C |  |
| 11 | ND |  | 1'' | 166.4, C |  |
| 12 | ND |  | 2'' | 55.5, CH | 4.69, dd (6.6, 2.7) |
| 13 | 15.1, CH<sub>3</sub> | 2.58, d (0.9) | NH-2'' |  | 7.64, d (6.6) |
| 14 | 7.7, CH<sub>3</sub> | 2.26, s | 3'' | 168.6, C |  |
| 1' | 166.1, C |  | 4'' | 58.6, CH | 3.57, m |
| 2' | 54.7, CH | 4.45, dd (6.4, 3.1) | NH-4'' |  | 7.75, m |
| NH-2' |  | 7.38, d (6.4) | 5'' | 31.8, CH | 2.10, m |
| 3' | 168.7, C |  | 6'' | 18.9, CH<sub>3</sub> | 1.10, d (6.7) |
| 4' | 57.9, CH | 3.56, m | 7'' | 19.2, CH<sub>3</sub> | 0.91, d (6.7) |
| NH-4' |  | 8.28, d (5.4) | 8'' | ND |  |
| 5' | 31.0, CH | 2.17, m | 9'' | 47.5, CH<sub>2</sub> | 3.82, m |
|  |  |  |  |  | 3.75, m |
| 6' | 18.8, CH<sub>3</sub> | 1.13, d (6.7) | 10'' | 22.8, CH<sub>2</sub> | 2.29, m |
|  |  |  |  |  | 2.07, m |
| 7' | 19.1, CH<sub>3</sub> | 0.87, d (6.7) | 11'' | 31.1, CH<sub>2</sub> | 2.84, m |
|  |  |  |  |  | 1.86, dd (11.9, 6.9) |
| 8' | ND |  | 12'' | 56.3, CH | 5.95, d (9.2) |
| 9' | 60.5, CH<sub>2</sub> | 4.49, d (13.4) | 13'' | 172.9, C |  |
|  |  | 4.07, d (13.4) |  |  |  |
| **10'** | 76.8, C |  | **14''** | 34.9, CH<sub>3</sub> | 2.89, s |
| **11'** | 43.1, CH<sub>2</sub> | 2.94, m | **15''** | 51.3, CH<sub>2</sub> | 4.72, d (17.1) |
|  |  | 2.25, m |  |  | 3.65, d (17.1) |
| **12'** | 56.7, CH | 6.27, d (10.5) | **16''** | 165.4, C |  |
| **13'** | 173.1, C |  | **17''** | 39.4, CH<sub>3</sub> | 2.93, s |
| **14'** | 35.1, CH<sub>3</sub> | 2.93, s | **18''** | 71.4, CH | 2.70, m |
| **15'** | 51.1, CH<sub>2</sub> | 4.36, d (17.1) | **19''** | 27.0, CH | 2.69, m |
|  |  | 3.58, d (17.1) |  |  |  |
| **16'** | 166.3, C |  | **20''** | 19.0, CH<sub>3</sub> | 0.75, d (6.3) |
| **17'** | 39.1, CH<sub>3</sub> | 2.95, s | **21''** | 21.7, CH<sub>3</sub> | 0.98, d (6.0) |
| **18'** | 71.3, CH | 2.67, m | **22''** | 167.2, C |  |
| **19'** | 26.9, CH | 2.65, m | **23''** | 75.2, CH | 5.22, qd (6.3, 2.7) |
| **20'** | 19.0, CH<sub>3</sub> | 0.75, d (6.8) | **24''** | 17.9, CH<sub>3</sub> | 1.30, d (6.3) |
| **21'** | 21.6, CH<sub>3</sub> | 0.96, d (6.4) |  |  |  |

**Table 2.** Antibacterial activities of compounds expressed as Minimal Inhibitory Concentrations (MIC)

| Strain | Actinomycin X<sub>2</sub> | Actinomycin D | Actinomycin L<sub>1</sub> | Actinomycin L<sub>2</sub> |
| --- | --- | --- | --- | --- |
| *S. aureus* MB5393<br>(methicillin-resistant) | 2-4 # | 1-2 | 4-8 | 8-16 |
| *S. aureus* ATCC29213<br>(methicillin-sensitive) | 1-2 | 1-2 | 2-4 | n.a. |
| *E. faecium* (vancomycin-sensitive)* | 1-2 | 1-2 | 4-8 | 8-16 |
| *E. faecium* VanB<br>(vancomycin-resistant)* | 1-2 | 1-2 | 4-8 | 8-16 |
| *S. epidermidis* * | 2-4 | 2-4 | 4-8 | n.a. |
| *E. coli* ATCC25922 | >128 | >128 | >128 | n.a |
| *K. pneumoniae* ATCC700603 | >128 | >128 | >128 | n.a |

(\*), clinical isolates; n.a.: data not available. # all concentrations given in µg/ml.

### Antimicrobial activity assay and MIC determination

The antimicrobial activity of the compounds was tested in liquid inhibition assays against seven pathogens including Gram-negative and Gram-positive bacteria (*Escherichia coli* ATCC25922, *Klebsiella pneumoniae* ATCC700603, methicillin-resistant *Staphylococcus aureus* MB5393, methicillin-sensitive *Staphylococcus aureus* ATCC29213, linezolid-resistant *Staphylococcus. epidermidis* (clinical isolate), vancomycin-sensitive *Enterococcus. faecium* (clinical isolate), and vancomycin-resistant *Enterococcus faecium* VanB (clinical isolate), as described (1). Each compound was serially diluted in DMSO with a dilution factor of 2 to test 10 concentrations starting at 128 μg/mL in all the antimicrobial assays. The MIC was defined as the lowest concentration of compound that inhibited ≥95% of the growth of a microorganism after overnight incubation. The Genedata Screener software (Genedata, Inc., Basel, Switzerland) was used to process and analyze the data and to calculate the RZ’ factor in the assay that was between 0.90 and 0.98 supporting its robustness.

### LC-MS/MS analysis

For LC-MS analyses, extracts were dissolved in MeOH to a final concentration of 1 mg/mL, and 1 μL was injected into Waters Acquity UPLC system equipped with Waters Acquity HSS C_18_ column (1.8 μm, 100 Å, 2.1 × 100 mm). For the LC, solvent A was 95% H_2_O, 5% acetonitrile (ACN) and 0.1% formic acid; solvent B was 100% ACN and 0.1% formic acid. The gradient used was 2% B for 1 min, 2–85% for 9 min, 85–100% for 1 min, and 100% for 3 min. The flow rate used was 0.5 mL/min. As for the MS, the following ESI source parameters were used: capillary voltage 3 kV, source temperature 325 °C, drying gas flow rate 10 L/min, and fragmentor 175 V. Full MS spectra were acquired in positive mode in the range of 100–1700 *m/z*, in the extended dynamic range mode. Internal reference masses of purine and HP-921 (Agilent) were continuously delivered to the ESI source through an Agilent 1260 isocratic pump.

LC-MS/MS acquisition was performed using Shimadzu Nexera X2 UHPLC system coupled to Shimadzu 9030 QTOF mass spectrometer, equipped with a standard ESI source unit, in which a calibrant delivery system (CDS) is installed. The dry extracts were dissolved in MeOH to a final concentration of 1 mg/mL, and 1 μL was injected into a Waters Acquity HSS C_18_ column (1.8 μm, 100 Å, 2.1 × 100 mm). The column was maintained at 40 °C, and run at a flow rate of 0.5 mL/min, using 0.1% formic acid in H_2_O as solvent A, and 0.1% formic acid in acetonitrile as solvent B. A gradient was employed for chromatographic separation starting at 5% B for 1 min, then 5–85% B for 9 min, 85–100% B for 1 min, and finally held at 100% B for 4 min. All the samples were analyzed in positive polarity, using data dependent acquisition mode. In this regard, full scan MS spectra (*m/z* 100–2000, scan rate 20 Hz) were followed by three data dependent MS/MS spectra (*m/z* 100–2000, scan rate 20 Hz) for the three most intense ions per scan using collision induced dissociation (CID) with collision energy ramp (CE 20–50 eV), and excluded for 0.05 s (one MS scan) before being re-selected for fragmentation. The parameters used for the ESI source were: interface voltage 4 kV, interface temperature 300 °C, nebulizing gas flow 3 L/min, and drying gas flow 10 L/min.

LC–MS/MS acquisition for molecular networking was performed using Thermo Instruments MS system (LTQ Orbitrap XL, Bremen, Germany) equipped with an electrospray ionization source (ESI). The Waters Acquity UPLC system equipped with Waters Acquity PDA was run using a SunFire Waters C_18_ column (3.5 μm, 100 Å, 4.6×150 mm), at a flow rate of 0.9 mL/min. Solvent A was 95% H_2_O, 5% acetonitrile (ACN) and 0.1% formic acid; solvent B was 100% ACN and 0.1% formic acid. The gradient used was 2% B for 1 min, 2–85% for 15 min, 85–100% for 3 min, and 100% for 3 min. As for the MS, the following ESI parameters were used: capillary voltage 5 V, spray voltage 3.5 kV, capillary temperature 300 °C, auxiliary gas flow rate 10 arbitrary units, and sheath gas flow rate 50 arbitrary units. Full MS spectra were acquired in the Orbitrap in positive mode at a mass range of 100–2000 *m/z*, and FT resolution of 30,000. Data-dependent MS^2^ spectra were acquired in the ion trap for the three most intense ions using collision induced dissociation (CID).

### Computation of mass spectral networks

MS/MS raw data were converted to a 32-bit mzXML file using MSConvert (ProteoWizard) (8) and spectral networks were assembled using Global Natural Product Social molecular networking (GNPS) (https://gnps.ucsd.edu) as described (53). The precursor ion mass tolerance was set to 2.0 Da and a MS/MS fragment ion tolerance of 0.5 Da, while the minimum cosine score was set to 0.7. The data were clustered using MSCluster with a minimum cluster size of three spectra. The spectra in the network were also searched against GNPS spectral libraries. A minimum score of 0.7 was set for spectral library search, with at least six fragment peaks matching. Cytoscape 3.7.2 was used for visualization of the generated molecular networks (45). The edge thickness was set to represent the cosine score, with thicker lines indicating higher similarity between nodes. LC– MS/MS data were deposited in the MassIVE Public GNPS data set (MSV000085106). The molecular networking job in GNPS can be found at https://gnps.ucsd.edu/ProteoSAFe/status.jsp?task=0c9153470404488d8927289139f875d3. The annotated MS/MS spectra were deposited in the GNPS spectral library for actinomycin L_1_ (CCMSLIB00005718892) and L_2_ (CCMSLIB00005718891).

### Statistical analysis

Prior to statistical analysis, mzXML files, which were converted using Shimadzu LabSolutions Postrun Analysis, were imported into Mzmine 2.31 (36) and processed as previously described (32). The aligned peak list was exported as a comma-separated file for statistical analysis. Statistical analysis was performed using MetaboAnalyst (9), where log transformation and pareto scaling was initially applied to the data. The normalized data were subjected to principal components analysis (PCA) and orthogonal partial least squares discriminant analysis (OPLS-DA). The quality of the models was evaluated with the relevant *R*^*2*^ and *Q*^*2*^. To identify the difference in intensity of a single mass feature among multiple growth conditions, one-way ANOVA was performed, followed by a *post hoc* Tukey’s honest significant difference (HSD) test.

## Supporting information

All Supplemental Information

## ACKNOWLEDGEMENTS

We are grateful to Victor Carrion for discussions. The work was funded by the NACTAR program of the Netherlands Organization for Scientific Research (NWO), grant nr. 16440. The authors declare no conflict of interests.

## Supplemental Tables and Figures

**Table S1**. X-ray Crystallography data.

**Table S2**. Accurate mass of [M+H]^+^ and product ions of the PPLs analyzed be ESI-QTOF MS/MS.

**Table S3**. Gene organization of the actinomycins biosynthetic gene cluster in *Streptomyces* sp. MBT27 and similarities to corresponding protein sequences encoded by orthologues in the *S. antibioticus* IMRU 3720 BGC.

**Fig. S1** Antimicrobial activity of *Streptomyces* sp. MBT27 extracts against *B. subtilis*.

**Fig. S2** Permutation validation of OPLS-DA model.

**Fig. S3** HRMS spectrum of **1.**

**Fig. S4** HRMS spectrum of **2**.

**Fig. S5** ^1^H NMR spectrum of **1** (850 MHz, in CDCl_3_ with TMS).

**Fig. S6** ^1^H–^1^H TOCSY spectrum of **1** (850 MHz, in CDCl_3_ with TMS).

**Fig. S7** ^1^H–^1^H COSY spectrum of **1** (850 MHz, in CDCl_3_ with TMS).

**Fig. S8** HSQC spectrum of **1** (850 MHz, in CDCl_3_ with TMS).

**Fig. S9** HMBC spectrum of **1** (850 MHz, in CDCl_3_ with TMS).

**Fig. S10** NOESY spectrum of **1** (850 MHz, in CDCl_3_ with TMS).

**Fig. S11** Stacked ^1^H NMR spectra of **1** and **2** (850 MHz, in CDCl_3_ with TMS).

**Fig. S12** UV spectrum of **1.**

**Fig. S13** UV spectrum of **2.**

**Fig. S14** IR spectrum of **1.**

**Fig. S15** IR spectrum of **2**

**Fig. S16** *S. antibioticus* is incapable to produces actinomycin L unless anthranilamide is supplied.

**Fig. S17** QTOF MS/MS spectrum of PPL 0.

**Fig. S18** QTOF MS/MS spectrum of PPL 1.

**Fig. S19** QTOF MS/MS spectrum of PPL 2.

**Fig. S20** QTOF MS/MS spectrum of PPL 3.

**Fig. S21 Alignment of the actinomycin BGCs from *S. antibioticus* and *Streptomyces* sp. MBT27**.

