## Supplementary material for "New from old: discovery of the novel antibiotic actinomycin L in *Streptomyces* sp. MBT27": All Supplemental Information

belonging to the manuscript

### SUPPLEMENTAL METHODS

#### Single Crystal X-ray Crystallography

All reflection intensities were measured at 100(2) K using a SuperNova diffractometer (equipped with Atlas detector) with Cu *K*α radiation (λ = 1.54178 Å) under the program CrysAlisPro (Version CrysAlisPro 1.171.39.29c, Rigaku OD, 2017). The same program was used to refine the cell dimensions and for data reduction. The structure was solved and refined with SHELXL-2018/3 (3). Analytical numeric absorption correction using a multifaceted crystal model was applied using CrysAlisPro. The temperature of the data collection was controlled using the system Cryojet (manufactured by Oxford Instruments). Prior to mounting the chosen single crystal on the diffractometer, crystals were placed in some Parabar 10312 on a microscope slide and cooled via a cold $N_2(g)$ stream in order to protect the crystal from potential solvent loss. The H atoms were placed at calculated positions (unless otherwise specified) using the instructions AFIX 13, AFIX 23, AFIX 43, AFIX 137 or AFIX 147 with isotropic displacement parameters having values 1.2 or 1.5 $U_{eq}$ of the attached C or O atoms. The H atoms attached to N2, N1', N2', N6', N7', N1" and N2" were found from difference Fourier maps, and their coordinates were refined pseudofreely using the DFIX instruction in order to keep the N–H bonds within an acceptable range. The structure is partly disordered.

The asymmetric unit contains one molecule of the target compound as well as some significant amount of lattice MeOH solvent molecules. Six solvent molecules were modelled as ordered with four being fully occupied and two with partial occupancy factors of 0.918(11) and 0.487(10). The contribution of the remaining amount of disordered lattice solvent molecules has been removed using the SQUEEZE procedure in Platon (4). Furthermore, the atoms C9", C10" and C11" are disordered over two orientations, and the occupancy factor of the major component of the disorder refines to 0.533(10). The absolute configuration has been established by anomalous-dispersion effects in diffraction measurements on the crystal, and the Flack and Hooft parameters refine to 0.08(5) and 0.06(5), respectively. The model has chirality S, S, R, R, R, S, S, S, R, R at C2', C2", C4', C4", C10'. C12', C18', C18", C23', C23", respectively.

### SUPPLEMENTAL TABLES AND FIGURES

**Table S1.** X-ray crystallography data.

| | xs2313a |
|---|---|
| Crystal data | |
| Chemical formula | $C_{69}H_{90}N_{14}O_{17}$·5.405($CH_4O$) |
| $M_r$ | 1560.87 |
| Crystal system, space group | Trigonal, $P3_221$ |
| Temperature (K) | 100 |
| $a$, $c$ (Å) | 18.4731 (2), 42.4216 (5) |
| $V$ (Å$^3$) | 12537.1 (3) |
| $Z$ | 6 |
| Radiation type | Cu $K\alpha$ |
| μ (mm$^{-1}$) | 0.77 |
| Crystal size (mm) | 0.23 × 0.15 × 0.07 |
| Data collection | |
| Diffractometer | SuperNova, Dual, Cu at zero, Atlas |
| Absorption correction | Analytical *CrysAlis PRO* 1.171.40.53 (Rigaku Oxford Diffraction, 2019) Analytical numeric absorption correction using a multifaceted crystal model based on expressions derived by (1). Empirical absorption correction using spherical harmonics, implemented in SCALE3 ABSPACK scaling algorithm. |
| $T_{min}$, $T_{max}$ | 0.897, 0.957 |
| No. of measured, independent and observed [$I > 2\sigma(I)$] reflections | 90598, 14994, 13477 |
| $R_{int}$ | 0.043 |
| (sin θ/λ)$_{max}$ (Å$^{-1}$) | 0.598 |
| Refinement | |
| $R[F^2 > 2\sigma(F^2)]$, $wR(F^2)$, $S$ | 0.046, 0.128, 1.03 |
| No. of reflections | 14994 |
| No. of parameters | 1091 |
| No. of restraints | 79 |

| | |
|---|---|
| H-atom treatment | H atoms treated by a mixture of independent and constrained refinement |
| $\Delta\rho_{max}$, $\Delta\rho_{min}$ (e Å$^{-3}$) | 0.68, -0.26 |
| Absolute structure | Flack x determined using 5493 quotients [(I+)-(I-)]/[(I+)+(I-)] (2). |
| Absolute structure parameter | 0.08 (5) |

**Table S2.** Accurate mass of $[M+H]^+$ and product ions of the PPLs analyzed be ESI-QTOF MS/MS.

| Compound | Formula | Proposed ion | Measurement (*m/z*) | Calculated (*m/z*) | Diff (ppm) |
|---|---|---|---|---|---|
| **PPL0** | $C_{31}H_{46}N_5O_9$ | $[M+H]^+$ | 632.3170 | 632.3295 | -18.99 |
| | $C_{31}H_{44}N_5O_8$ | $[M+H-H_2O]^+$ | 614.3177 | 614.3189 | -1.2 |
| | $C_{25}H_{33}N_4O_7$ | $[M+H-(H\text{-}MeVal\text{-}OH)]^+$ | 501.2347 | 501.2349 | 0.65 |
| | $C_{22}H_{28}N_3O_6$ | $[M+H-(H\text{-}Sar\text{-}MeVal\text{-}OH)]^+$ | 430.1969 | 430.1978 | -0.84 |
| | $C_{19}H_{35}N_4O_6$ | $[(H\text{-}Val\text{-}HyPro\text{-}Sar\text{-}MeVal\text{-}OH)+H]^+$ | 415.2552 | 415.2556 | 0.214 |
| | $C_{19}H_{33}N_4O_5$ | $[(H\text{-}Val\text{-}HyPro\text{-}Sar\text{-}MeVal\text{-}OH)+H-H_2O]^+$ | 397.2441 | 397.2450 | -1.124 |
| | $C_{14}H_{26}N_3O_5$ | $[(H\text{-}Hyp\text{-}Sar\text{-}MeVal\text{-}OH)+H]^+$ | 316.1870 | 316.1872 | 0.957 |
| | $C_{14}H_{24}N_3O_4$ | $[(H\text{-}Hyp\text{-}Sar\text{-}MeVal\text{-}OH)+H-H_2O]^+$ | 298.1761 | 298.1766 | -0.110 |
| | $C_{13}H_{22}N_3O_4$ | $[(H\text{-}Val\text{-}HyPro\text{-}Sar)+H]^+$ | 284.1606 | 284.1610 | 0.413 |
| | $C_{12}H_{12}NO_3$ | $[M+H-(H\text{-}Val\text{-}HyPro\text{-}Sar\text{-}MeVal\text{-}OH)]^+$ | 218.0816 | 218.0817 | 1.973 |
| | $C_9H_{19}N_2O_3$ | $[(H\text{-}Sar\text{-}MeVal\text{-}OH)+H]^+$ | 203.1391 | 203.1395 | 0.399 |
| | $C_6H_{14}NO_2$ | $[(H\text{-}MeVal\text{-}OH)+H]^+$ | 132.1015 | 132.1024 | -3.067 |
| | $C_4H_6NO_2$ | $[Thr+H]^+$ | 100.0389 | 100.0398 | -4.048 |
| **PPL1** | $C_{31}H_{46}N_5O_8$ | $[M+H]^+$ | 616.3326 | 616.3346 | -2.417 |
| | $C_{31}H_{44}N_5O_7$ | $[M+H-H_2O]^+$ | 598.3248 | 598.3241 | 2.131 |
| | $C_{25}H_{33}N_4O_6$ | $[M+H-(H\text{-}MeVal\text{-}OH)]^+$ | 485.2392 | 485.2400 | -0.538 |
| | $C_{22}H_{28}N_3O_5$ | $[M+H-(H\text{-}Sar\text{-}MeVal\text{-}OH)]^+$ | 414.2025 | 414.2028 | 0.368 |
| | $C_{19}H_{35}N_4O_5$ | $[(H\text{-}Val\text{-}Pro\text{-}Sar\text{-}MeVal\text{-}OH)+H]^+$ | 399.2607 | 399.2607 | 1.261 |
| | $C_{19}H_{33}N_4O_4$ | $[(H\text{-}Val\text{-}Pro\text{-}Sar\text{-}MeVal\text{-}OH)+H-H_2O]^+$ | 381.2504 | 381.2501 | 2.014 |
| | $C_{17}H_{21}N_2O_4$ | $[M+H-(H\text{-}Pro\text{-}Sar\text{-}MeVal\text{-}OH)]^+$ | 317.1496 | 317.1501 | 0.052 |
| | $C_{14}H_{26}N_3O_4$ | $[(H\text{-}Pro\text{-}Sar\text{-}MeVal\text{-}OH)+H]^+$ | 300.1916 | 300.1923 | -0.609 |
| | $C_{14}H_{24}N_3O_3$ | $[(H\text{-}Pro\text{-}Sar\text{-}MeVal\text{-}OH)+H-H_2O]^+$ | 282.1809 | 282.1817 | -1.127 |
| | $C_{13}H_{22}N_3O_3$ | $[(H\text{-}Val\text{-}Pro\text{-}Sar)+H]^+$ | 268.1657 | 268.1661 | 0.492 |
| | $C_9H_{19}N_2O_3$ | $[(H\text{-}Sar\text{-}MeVal\text{-}OH)+H]^+$ | 203.1392 | 203.1395 | 0.891 |
| | $C_8H_{13}N_2O_2$ | $[(H\text{-}Pro\text{-}Sar)+H]^+$ | 169.0968 | 169.0977 | -2.095 |
| | $C_6H_{14}NO_2$ | $[(H\text{-}MeVal\text{-}OH)+H]^+$ | 132.1016 | 132.1024 | -2.310 |
| | $C_4H_6NO_2$ | $[Thr+H]^+$ | 100.0391 | 100.0398 | -2.049 |
| **PPL2** | $C_{31}H_{44}N_5O_9$ | $[M+H]^+$ | 630.3132 | 630.3139 | -0.245 |
| | $C_{31}H_{42}N_5O_8$ | $[M+H-H_2O]^+$ | 612.3027 | 612.3033 | -0.146 |
| | $C_{25}H_{33}N_4O_8$ | $[M+H-(H\text{-}MeVal)]^+$ | 517.2286 | 517.2298 | -1.335 |
| | $C_{25}H_{31}N_4O_7$ | $[M+H-(H\text{-}MeVal\text{-}OH)]^+$ | 499.2190 | 499.2192 | 0.549 |
| | $C_{22}H_{26}N_3O_6$ | $[M+H-(H\text{-}Sar\text{-}MeVal\text{-}OH)]^+$ | 428.1814 | 428.1821 | -0.495 |
| | $C_{19}H_{33}N_4O_6$ | $[(H\text{-}Val\text{-}OxoPro\text{-}Sar\text{-}MeVal\text{-}OH)+H]^+$ | 413.2395 | 413.2400 | 0.094 |
| | $C_{19}H_{31}N_4O_5$ | $[(H\text{-}Val\text{-}OxoPro\text{-}Sar\text{-}MeVal\text{-}OH)+H-H_2O]^+$ | 395.2293 | 395.2294 | 1.021 |
| | $C_{14}H_{24}N_3O_5$ | $[(H\text{-}OxoPro\text{-}Sar\text{-}MeVal\text{-}OH)+H]^+$ | 314.1711 | 314.1715 | 0.168 |
| | $C_{14}H_{22}N_3O_4$ | $[(H\text{-}Val\text{-}OxoPro\text{-}Sar\text{-}MeVal\text{-}OH)+H-H_2O]^+$ | 296.1605 | 296.1610 | 0.059 |
| | $C_{13}H_{20}N_3O_4$ | $[(H\text{-}Val\text{-}OxoPro\text{-}Sar)+H]^+$ | 282.1448 | 282.1453 | -0.116 |
| | $C_{12}H_{12}NO_3$ | $[M+H-(H\text{-}Val\text{-}OxoPro\text{-}Sar\text{-}MeVal\text{-}OH)]^+$ | 218.0815 | 218.0817 | 1.514 |
| | $C_9H_{19}N_2O_3$ | $[(H\text{-}Sar\text{-}MeVal\text{-}OH)+H]^+$ | 203.1391 | 203.1395 | 0.399 |
| | $C_6H_{14}NO_2$ | $[(H\text{-}MeVal\text{-}OH)+H]^+$ | 132.1017 | 132.1024 | -1.554 |
| | $C_4H_6NO_2$ | $[Thr+H]^+$ | 100.0391 | 100.0398 | -2.049 |
| **PPL3** | $C_{38}H_{50}N_7O_9$ | $[M+H]^+$ | 748.3663 | 748.3670 | -0.204 |
| | $C_{38}H_{48}N_7O_8$ | $[M+H-H_2O]^+$ | 730.3561 | 730.3564 | 0.290 |
| | $C_{32}H_{37}N_6O_7$ | $[M+H-(H\text{-}MeVal\text{-}OH)]^+$ | 617.2716 | 617.2723 | -0.363 |
| | $C_{26}H_{39}N_6O_6$ | $[(H\text{-}Val\text{-}AntPro\text{-}Sar\text{-}MeVal\text{-}OH)+H]^+$ | 531.2928 | 531.2931 | 0.453 |
| | $C_{26}H_{37}N_6O_5$ | $[(H\text{-}Val\text{-}AntPro\text{-}Sar\text{-}MeVal\text{-}OH)+H-H_2O]^+$ | 513.2820 | 513.2825 | 0.010 |
| | $C_{21}H_{30}N_5O_5$ | $[(H\text{-}AntPro\text{-}Sar\text{-}MeVal\text{-}OH)+H]^+$ | 432.2247 | 432.2246 | 1.283 |
| | $C_{21}H_{28}N_5O_4$ | $[(H\text{-}AntPro\text{-}Sar\text{-}MeVal\text{-}OH)+H-H_2O]^+$ | 414.2137 | 414.2141 | 0.288 |
| | $C_{20}H_{26}N_5O_4$ | $[(H\text{-}Val\text{-}AntPro\text{-}Sar)+H]+$ | 400.1982 | 400.1984 | 0.673 |
| | $C_{17}H_{21}N_4O_3$ | $[(H\text{-}Val\text{-}AntPro)+H]+$ | 329.1607 | 329.1613 | -0.36 |
| | $C_{17}H_{21}N_2O_4$ | $[M+H-(H\text{-}AntPro\text{-}Sar\text{-}MeVal\text{-}OH)]^+$ | 317.1493 | 317.1501 | -0.894 |
| | $C_{15}H_{17}N_4O_3$ | $[(H\text{-}AntPro\text{-}Sar)+H]^+$ | 301.1295 | 301.1300 | -0.056 |
| | $C_9H_{19}N_2O_3$ | $[(H\text{-}Sar\text{-}MeVal\text{-}OH)+H]^+$ | 203.1390 | 203.1395 | -0.093 |
| | $C_6H_{14}NO_2$ | $[(H\text{-}MeVal\text{-}OH)+H]^+$ | 132.1014 | 132.1024 | -3.824 |
| | $C_4H_6NO_2$ | $[Thr+H]^+$ | 100.0401 | 100.0398 | 7.947 |

**Table S3.** Gene organization of the actinomycins biosynthetic gene cluster in *Streptomyces* sp. MBT27 and similarities to corresponding protein sequences encoded by orthologues in the *S. antibioticus IMRU 3720* biosynthetic gene cluster.

| ***S. antibioticus IMRU 3720*** **gene** | **Function** | ***Streptomyces*** **sp. MBT27 actinomycin cluster ORF** | **Identity** | **Similarity** |
|---|---|---|---|---|
| ***saacmT*** | Hypothetical protein | ORF01 | 0.80 | 0.89 |
| ***saacmS*** | Hypothetical protein | ORF02 | 0.84 | 0.93 |
| ***saacmR*** | MbtH-like protein | ORF03 | 0.86 | 0.94 |
| ***saacmD*** | 4-MHA carrier protein AcmACP | ORF04 | 0.60 | 0.69 |
| ***saacmA*** | Peptide synthetase ACMS I | ORF05 | 0.71 | 0.78 |
| ***saacmB*** | Peptide synthetase ACMS II | ORF06 | 0.70 | 0.78 |
| ***saacmC*** | Peptide synthetase ACMS III | ORF07 | 0.77 | 0.85 |
| ***saacmE*** | Hypothetical protein | ORF08 | 0.91 | 0.93 |
| ***saacmF*** | Aryl formamidase | ORF09 | 0.79 | 0.82 |
| ***saacmG*** | Tryptophan 2,3-dioxygenase | ORF10 | 0.83 | 0.89 |
| ***saacmK*** | Kynureninase | ORF11 | 0.86 | 0.91 |
| ***saacmL*** | Methyltransferase | ORF12 | 0.88 | 0.93 |
| ***saacmM*** | Cytochrome P450 | ORF13 | 0.85 | 0.88 |
| ***saacmN*** | Ferredoxin | ORF14 | 0.69 | 0.78 |
| ***saacmO*** | LmbU-like protein | ORF15 | 0.72 | 0.80 |
| ***saacmP*** | TetR family transcriptional regulator | ORF16 | 0.77 | 0.86 |
| ***saacmQ*** | Siderophore-interacting protein | ORF17 | 0.76 | 0.85 |
| ***saacmrA*** | ABC transporter ATPase subunit | ORF18 | 0.89 | 0.93 |
| ***saacmrB*** | ABC 2-type transporter | ORF19 | 0.93 | 0.96 |
| ***saacmrC*** | UvrA-like protein | ORF20 | 0.86 | 0.90 |

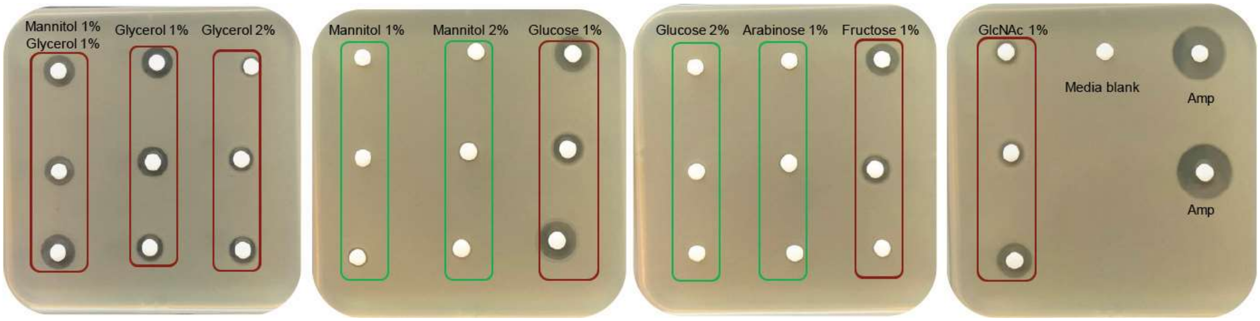

**Fig. S1 Representative results of antimicrobial activity of *Streptomyces* sp. MBT27 extracts against *B. subtilis.*** *Streptomyces* sp. MBT27 was fermented in minimal medium (MM) with different carbon sources, namely (percentages in w/v): 1% of both mannitol and glycerol, 1% mannitol, 2% mannitol, 1% glycerol, 2% glycerol, 1% glucose, 2% glucose, 1% fructose, 1% arabinose, or 1% *N*-acetylglucosamine (GlcNAc), and extracted with ethyl acetate. The red boxes indicate active extracts of *Streptomyces* sp. MBT27, while the green boxes represent extracts without antibacterial activity against *B. subtilis*. Ampicillin was used as a positive control. Note, that extracts of cultures fermented with different carbon sources showed different bioactivity profiles.

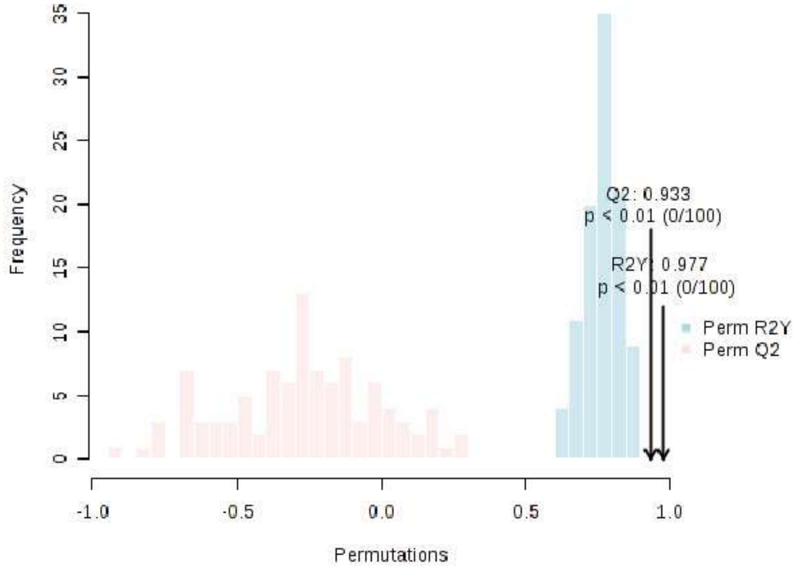

**Fig. S2** Permutation validation of OPLS-DA model.

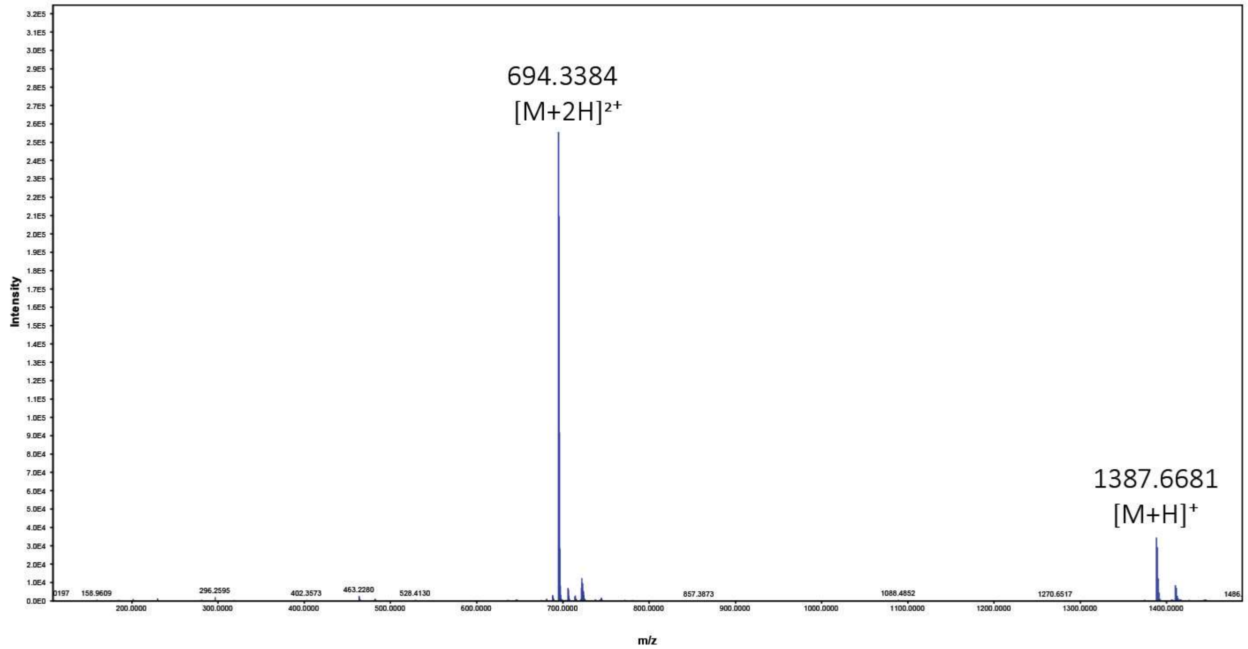

**Fig. S3** HRMS spectrum of **1.**

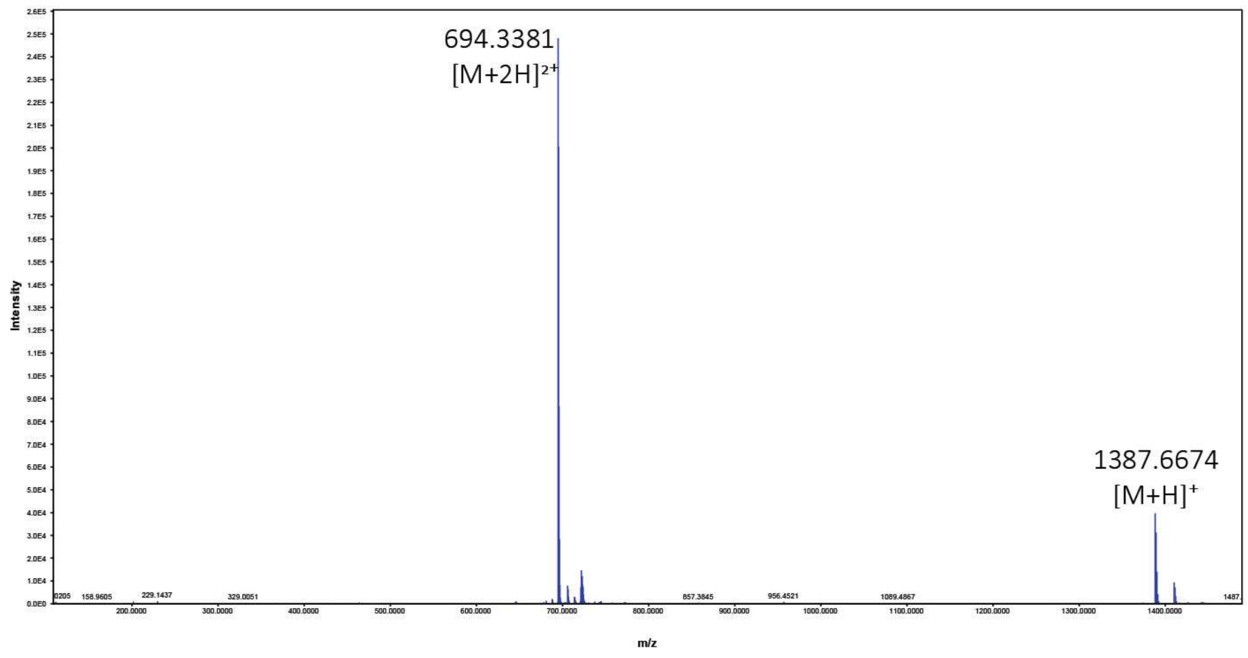

**Fig. S4** HRMS spectrum of **2.**

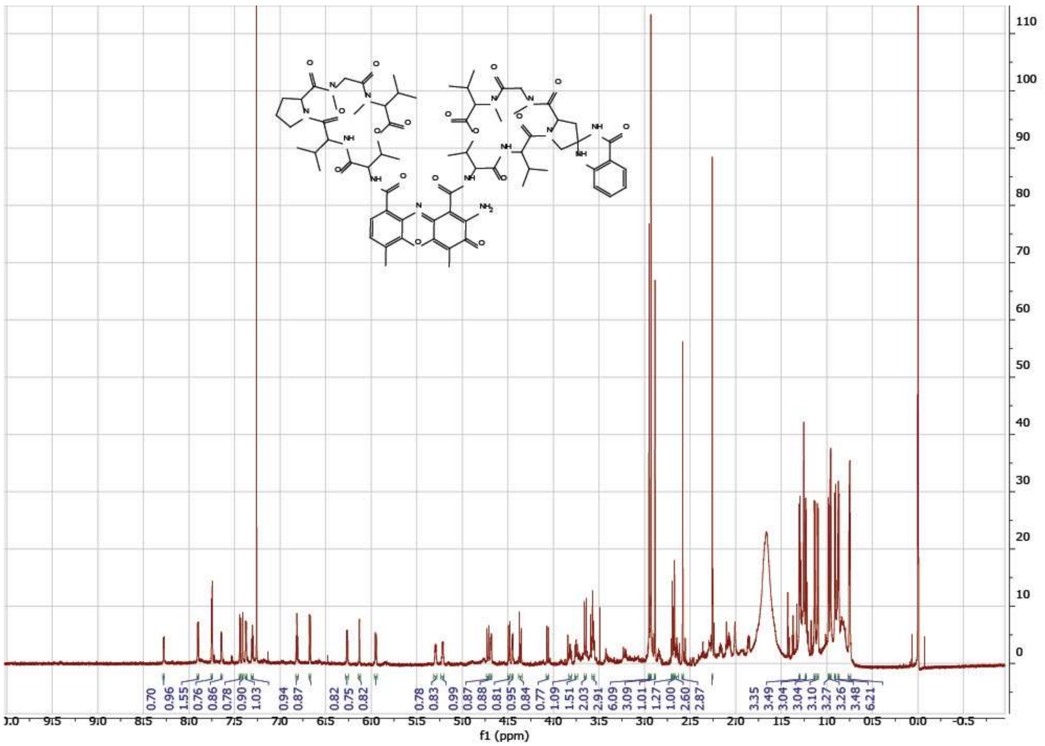

**Fig. S5** $^{1}$H NMR spectrum of **1** (850 MHz, in $CDCl_3$ with TMS).

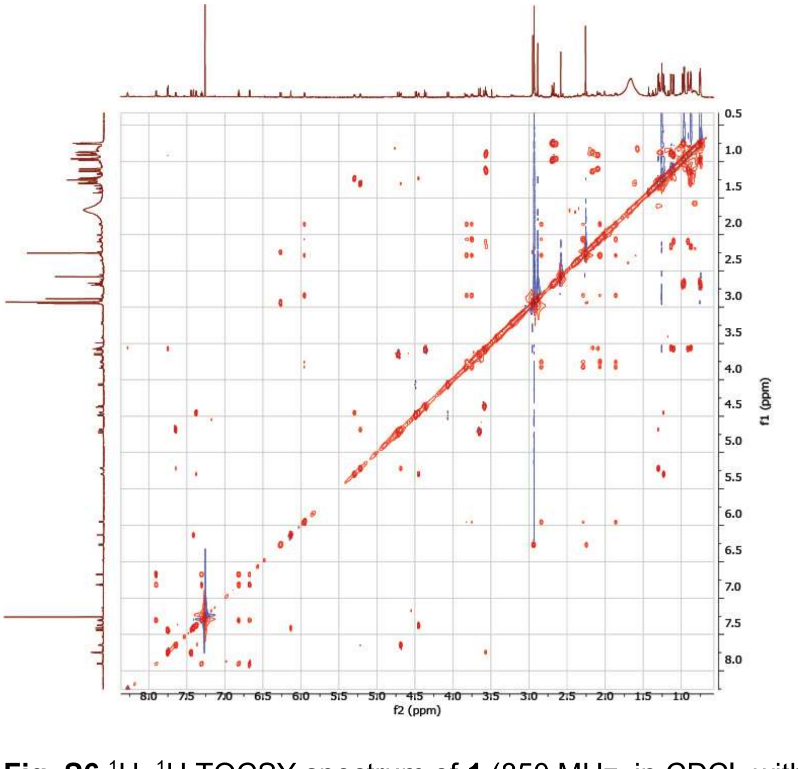

**Fig. S6** $^{1}$H–$^{1}$H TOCSY spectrum of **1** (850 MHz, in $CDCl_3$ with TMS).

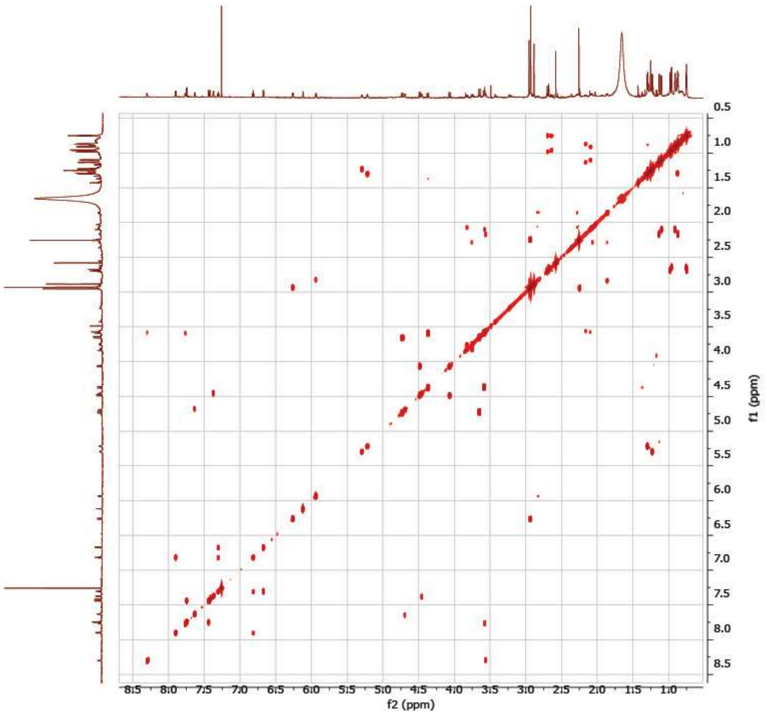

**Fig. S7** $^1H$–$^1H$ COSY spectrum of **1** (850 MHz, in $CDCl_3$ with TMS).

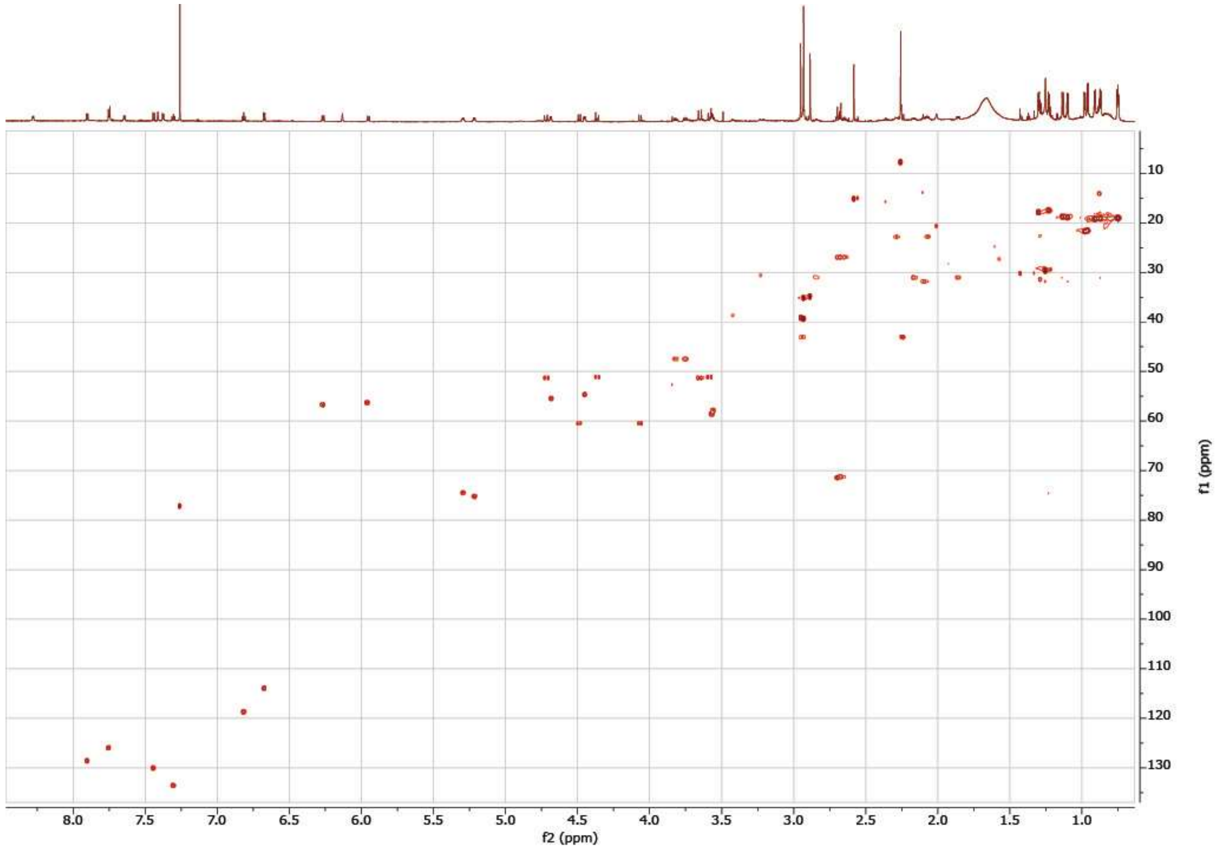

**Fig. S8** HSQC spectrum of **1** (850 MHz, in $CDCl_3$ with TMS).

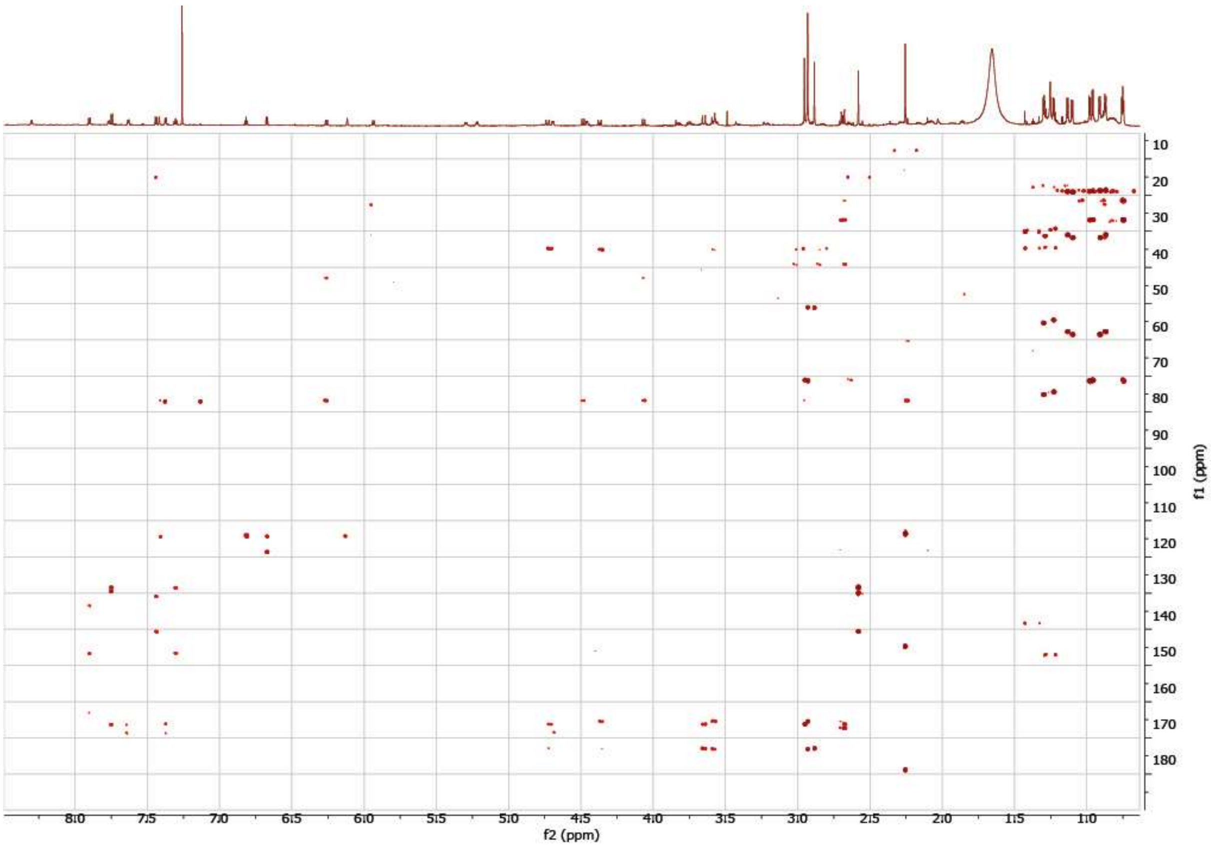

**Fig. S9** HMBC spectrum of **1** (850 MHz, in $CDCl_3$ with TMS).

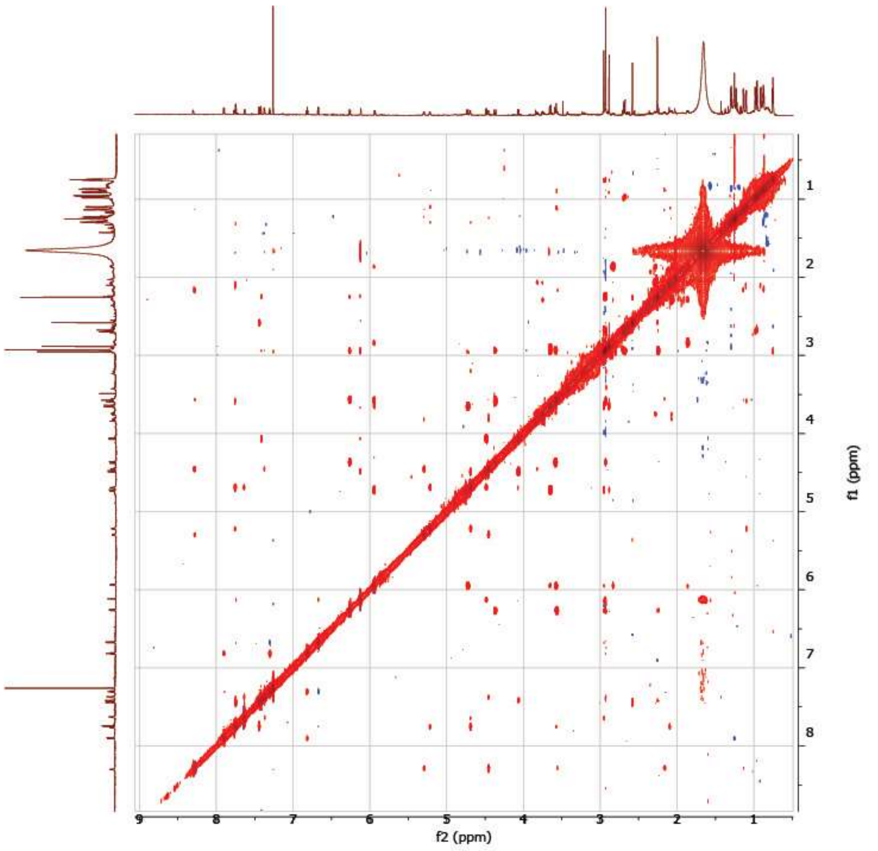

**Fig. S10** NOESY spectrum of **1** (850 MHz, in $CDCl_3$ with TMS).

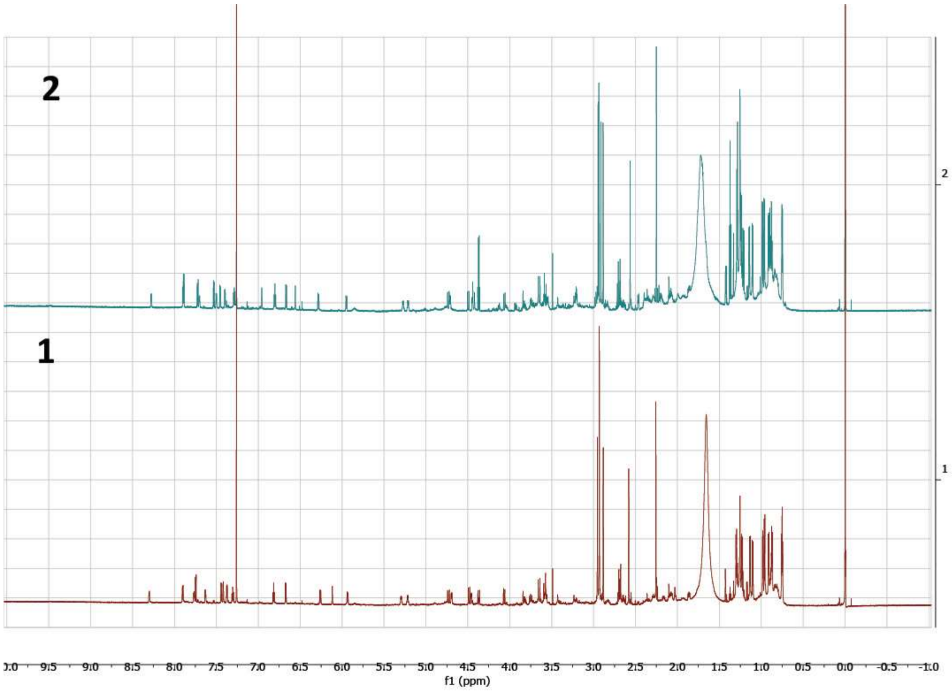

**Fig. S11** Stacked $^1$H NMR spectra of **1** and **2** (850 MHz, in $CDCl_3$ with TMS).

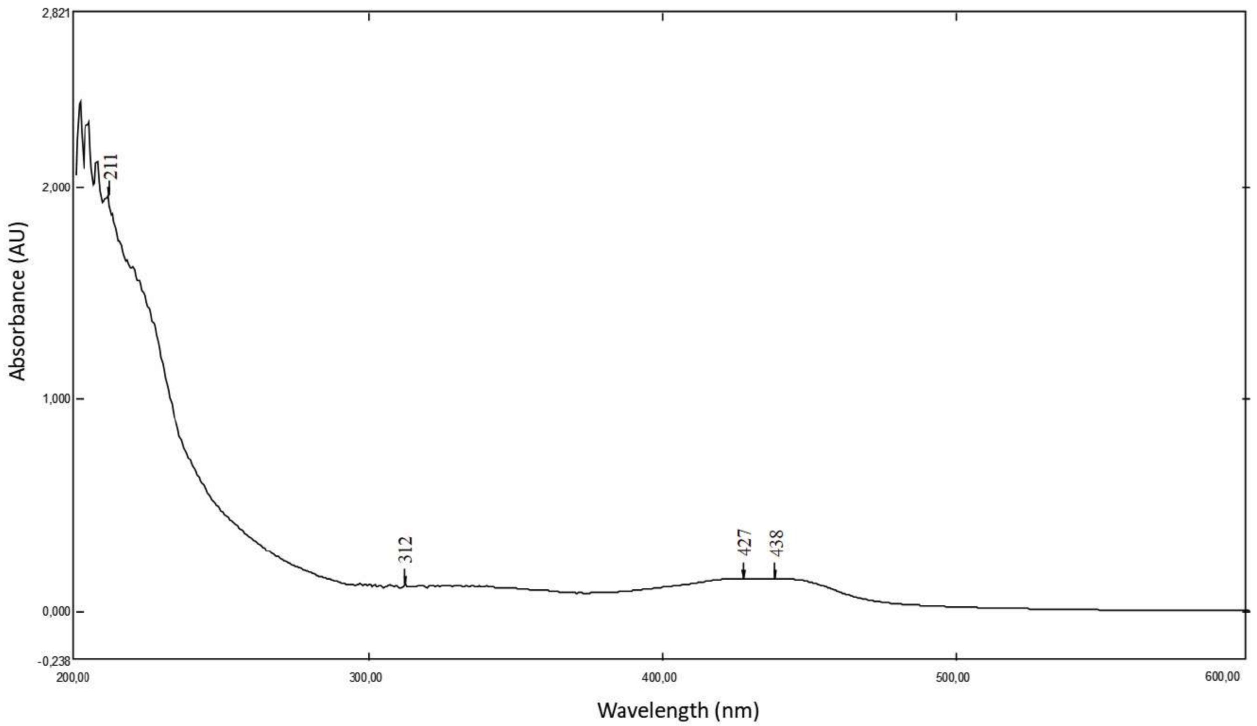

**Fig. S12** UV spectrum of **1.**

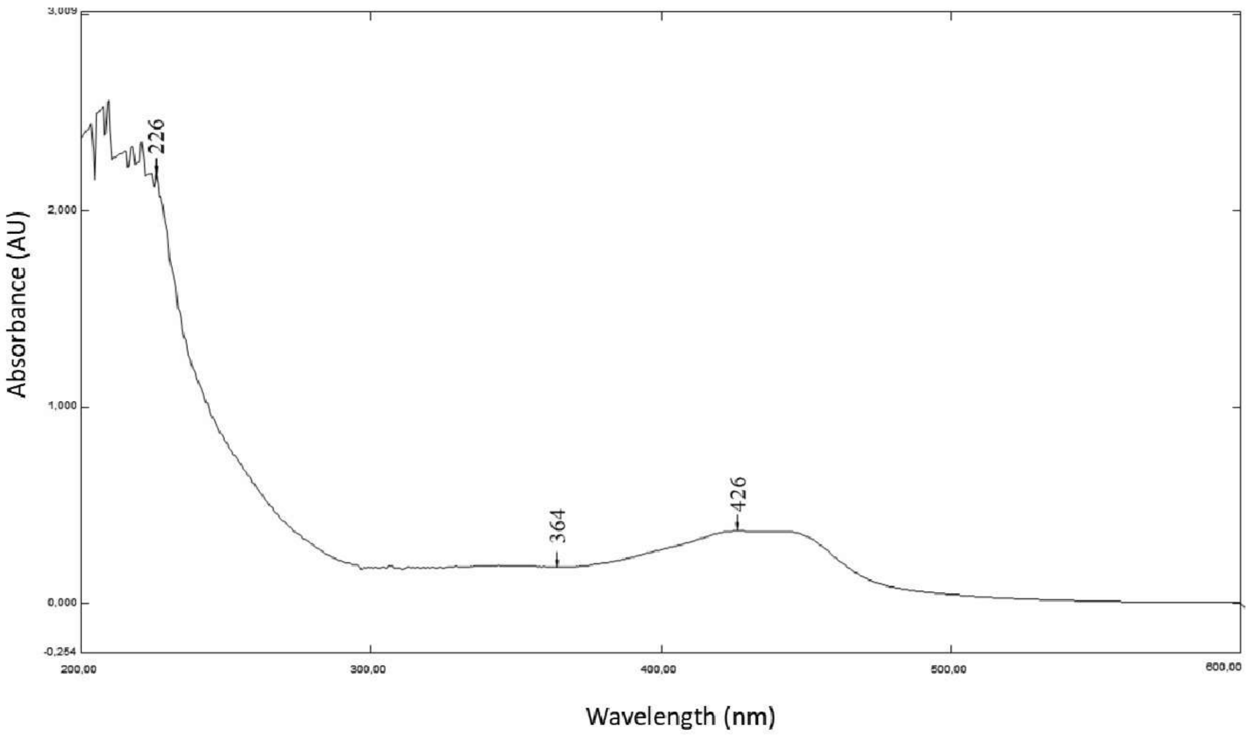

**Fig. S13** UV spectrum of **2.**

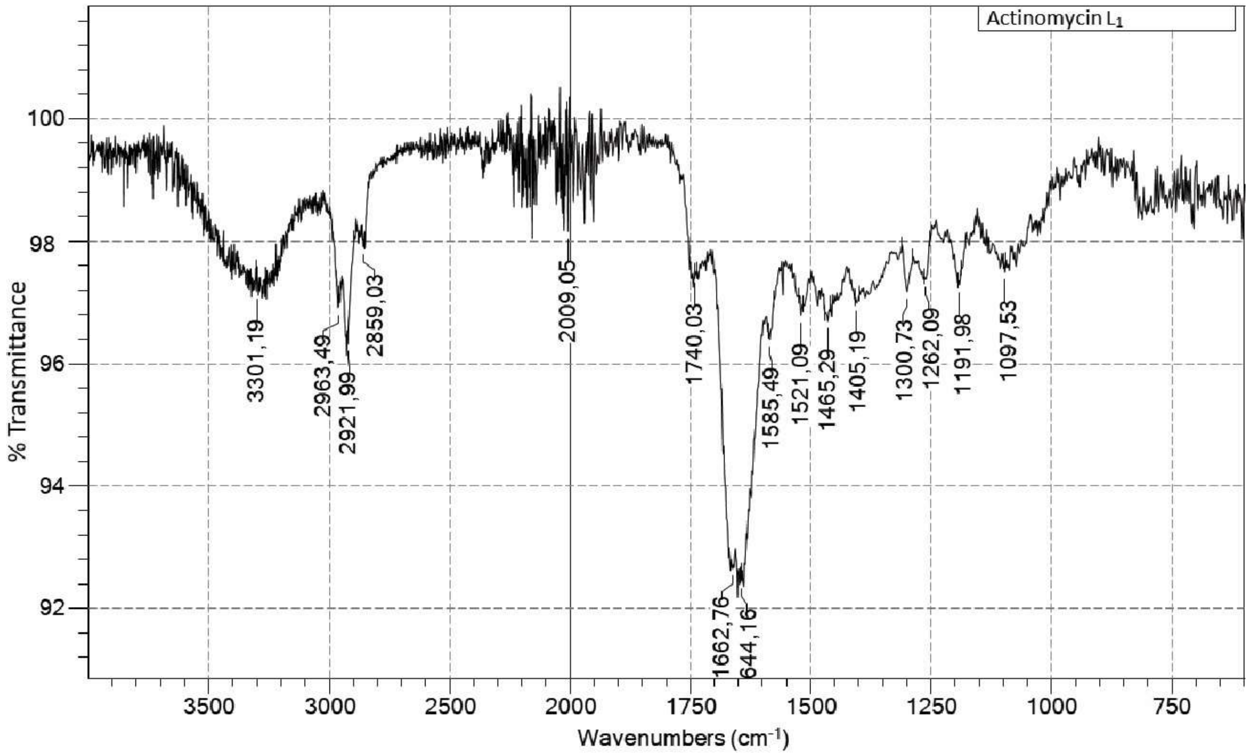

**Fig. S14** IR spectrum of **1.**

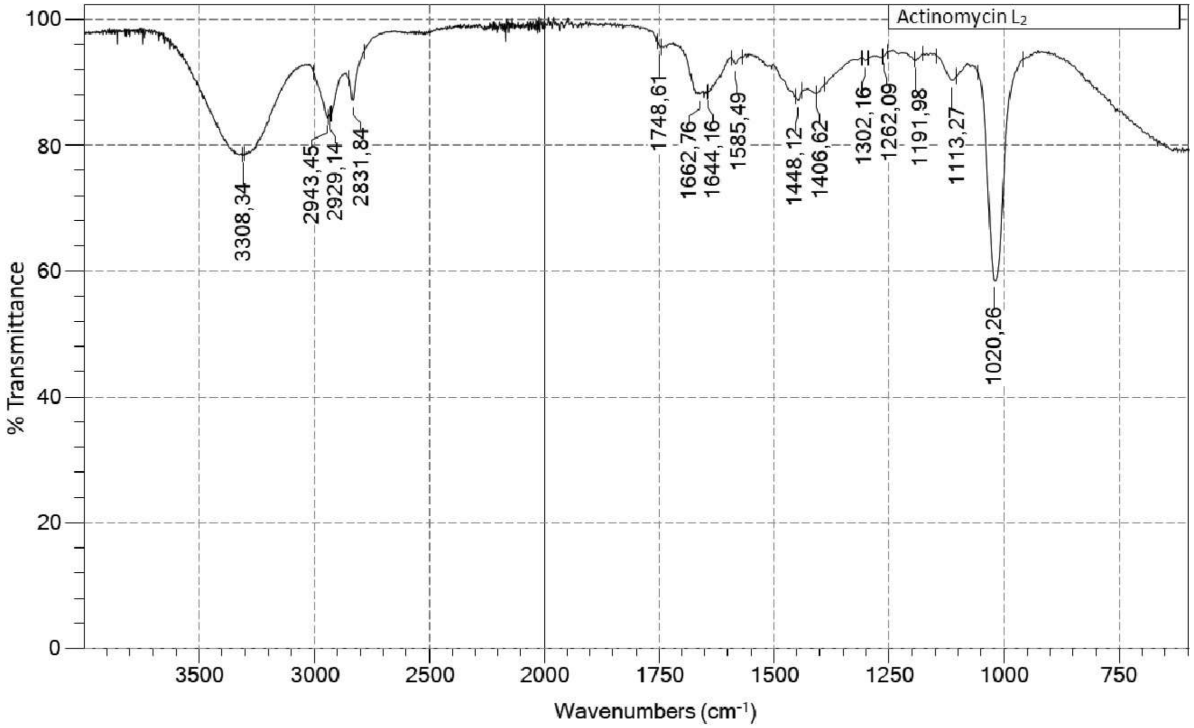

**Fig. S15** IR spectrum of **2.**

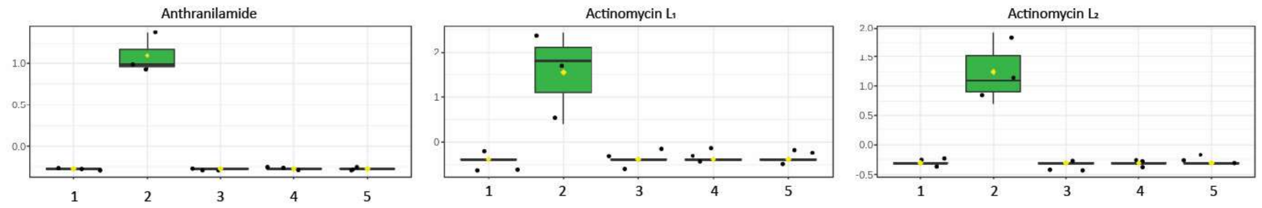

**Fig. S16** *S. antibioticus* is incapable to produces actinomycin L unless anthranilamide is supplied. Box plots showing the relative intensities of anthranilamide, actinomycin $L_1$ and $L_2$ in the cultures of *S. antibioticus* fermented for seven days in MM with different carbon sources 1. 1% fructose; 2. 1% fructose + 0.7 mM anthranilamide; 3. 1% glycerol; 4. 1% mannitol + 1 % glycerol; 5. 2% glycerol (all %ages in w/v).

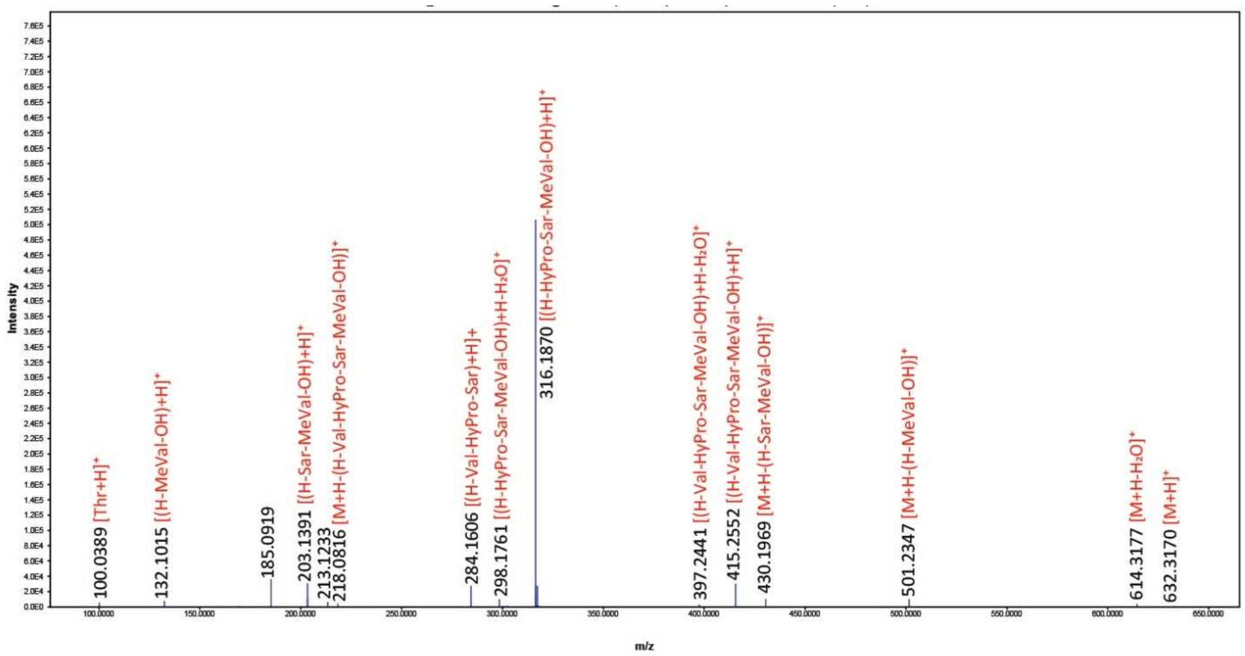

**Fig. S17** QTOF MS/MS spectrum of PPL 0.

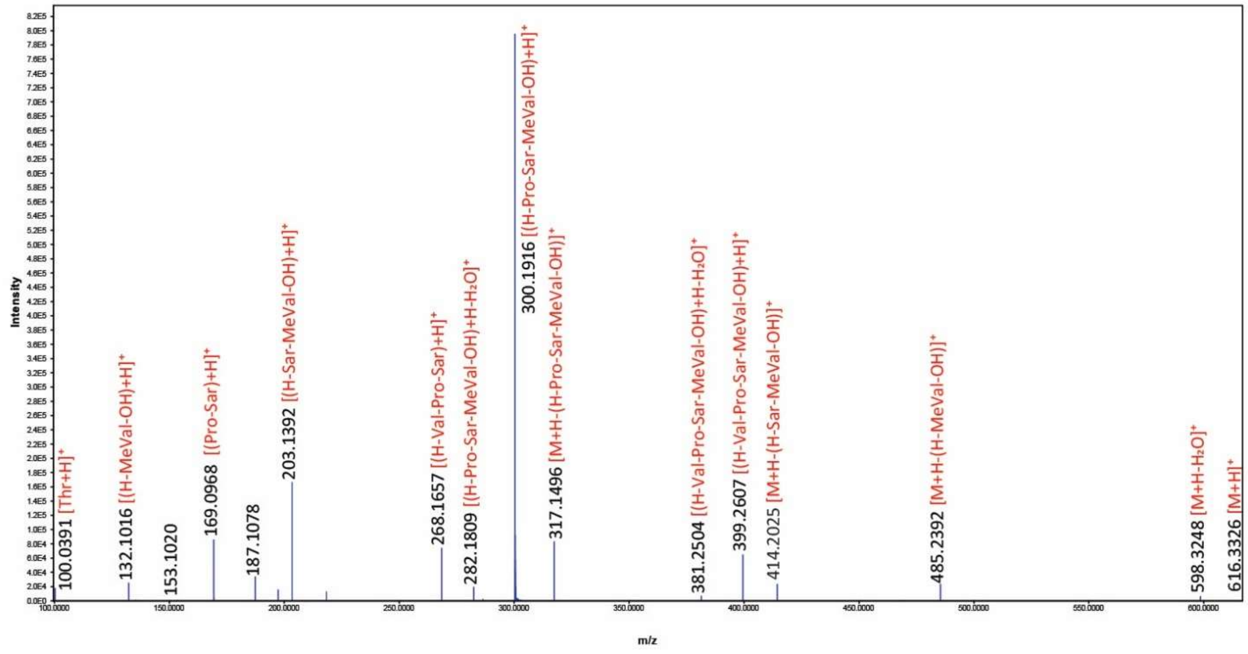

**Fig. S18** QTOF MS/MS spectrum of PPL 1.

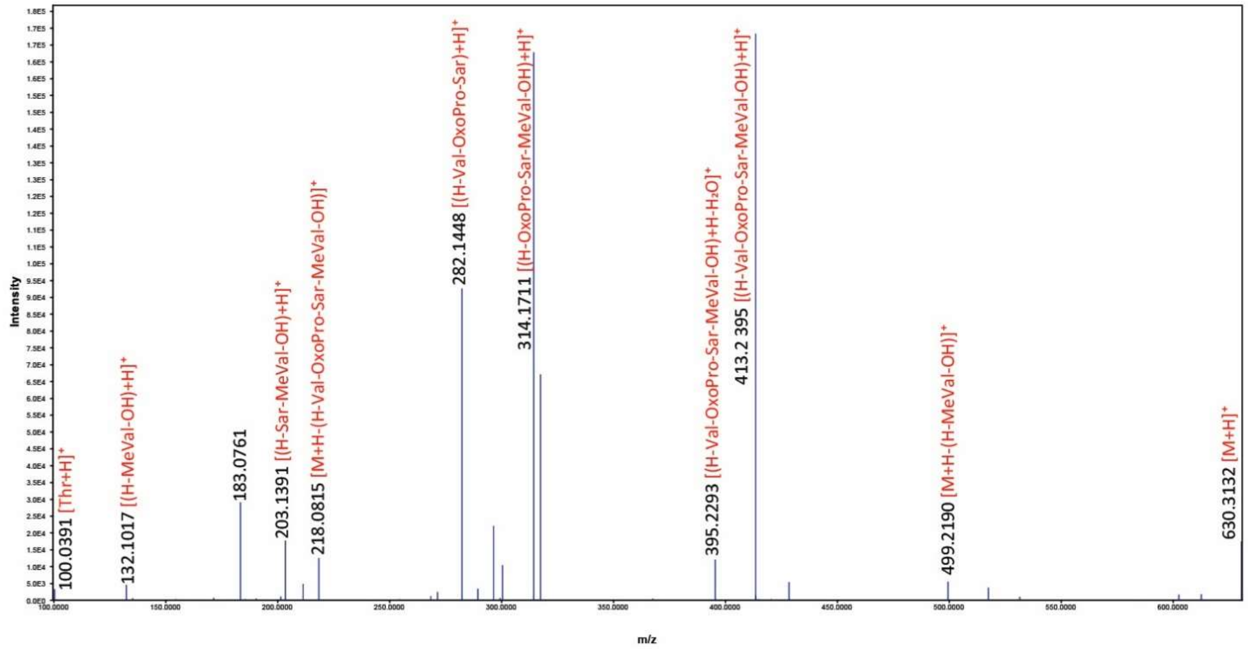

**Fig. S19** QTOF MS/MS spectrum of PPL 2.

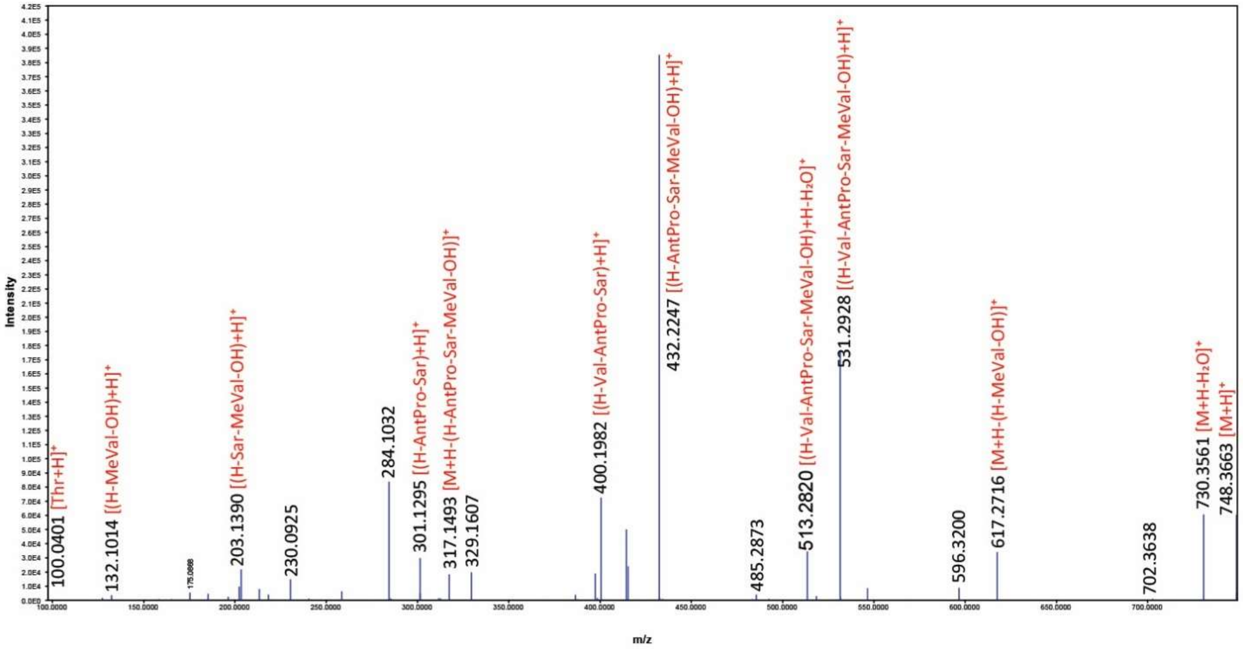

**Fig. S20** QTOF MS/MS spectrum of PPL 3.

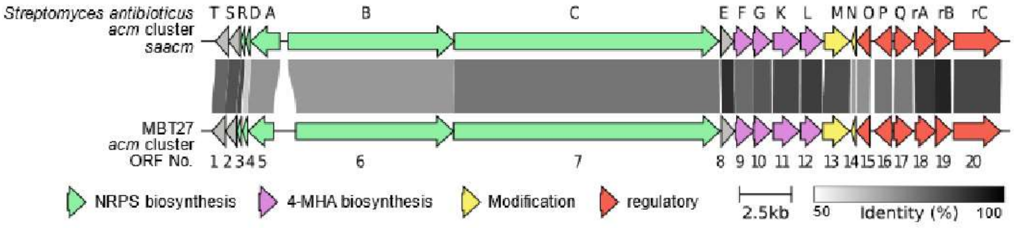

**Fig. S21 Alignment of actinomycin biosynthetic gene cluster from *S. antibioticus* and *Streptomyces* sp. MBT27.** Grey arrows indicate genes with unknown function. Grey bars connecting homologues pairs from the two cluster, the identity of two genes is indicated by different levels of greyness.
